# Hemin shows antiviral activity *in vitro*, possibly through suppression of viral entry mediators

**DOI:** 10.1101/2022.05.24.493187

**Authors:** Mehmet Altay Unal, Ceylan Verda Bitirim, Julia Somers, Gokce Yagmur Summak, Omur Besbinar, Ebru Kocakaya, Cansu Gurcan, Hasan Nazir, Zeynep Busra Aksoy Ozer, Sibel Aysil Ozkan, Sidar Bereketoglu, Aykut Ozkul, Emek Demir, Kamil Can Akcali, Acelya Yilmazer

## Abstract

Heme oxygenase-1 (HO-1) is a stress-induced enzyme that catalyzes the breakdown of heme into biliverdin, carbon monoxide, and iron. Targeting HO-1 to treat severe COVID-19 has been suggested by several groups, yet the role of HO-1 in SARS-CoV-2 infection remains unclear. Based on this, we aimed to investigate the antiviral activity of Hemin, an activator of HO-1. Infectivity of SARS-CoV-2 was decreased in Vero E6 cells treated with Hemin. Hemin also decreased TMPRSS2 and ACE2 mRNA levels in non-infected cells, possibly explaining the observed decrease in infectivity. TMPRSS2 protein expression and proteolytic activity were decreased in Vero E6 cells treated with Hemin. Besides that, experimental studies supported with in silico calculations. Overall, our study supports further exploration of Hemin as a potential antiviral and inflammatory drug for the treatment of COVID-19.

## Introduction

Although the development of effective vaccines against COVID-19 is encouraging, many unknowns remain with respect to the biology of this insidious disease. Even in light of recent advancements, understanding the molecular mechanisms that influence SARS-CoV-2 infection remains of critical importance. Due to worldwide urgency numerous efforts have been made to evaluate drug repurposing candidates (McKee et al., 2020). This line of work has the potential to identify valuable therapeutics but also contributes to our understanding of SARS-CoV-2 biology. Drug perturbation of infection models not only screens for agents which may benefit COVID-19 patients, but also gives valuable insight as to the molecular mechanisms governing the disease.

In light of this, we sought to evaluate the clinically approved drug Hemin^®^ (Ovation Pharmaceuticals, Inc., USA), also known as hemin, which is a known activator of the potent antiviral, antioxidant, and anti-inflammatory heme oxygenase-1 (HO-1) axis (X. Chen et al., 2018). Hemin is a heme analog consisting of a porphyrin ring and central iron (Figure 1A). HO-1 is a stress-induced enzyme that catalyzes the breakdown of the iron-containing porphyrin, heme, into biliverdin, carbon monoxide, and iron (B. Wu et al., 2019). Heme catabolism by HO-1 has demonstrated benefits in several conditions with strong relevance to COVID-19 including; acute lung injury (X. Chen et al., 2018), sepsis (Takaki et al., 2010), and ARDS (Nagasawa et al., 2020). Antiviral activity of HO-1 has also been reported in dengue virus, hepatitis C virus, influenza, and respiratory syncytial virus (Espinoza et al., 2017; Olagnier et al., 2014; Tseng et al., 2016; Wang et al., 2017; Zhu et al., 2008). This is partially because HO-1 and its associates counteract oxidative stress, which some viruses use to promote their life cycle. Additionally, HO-1 can activate the type-1 interferon response which is critical during early infection and is a key part of innate immunity (Lehmann et al., 2010; Tseng et al., 2016).

**Figure1:**
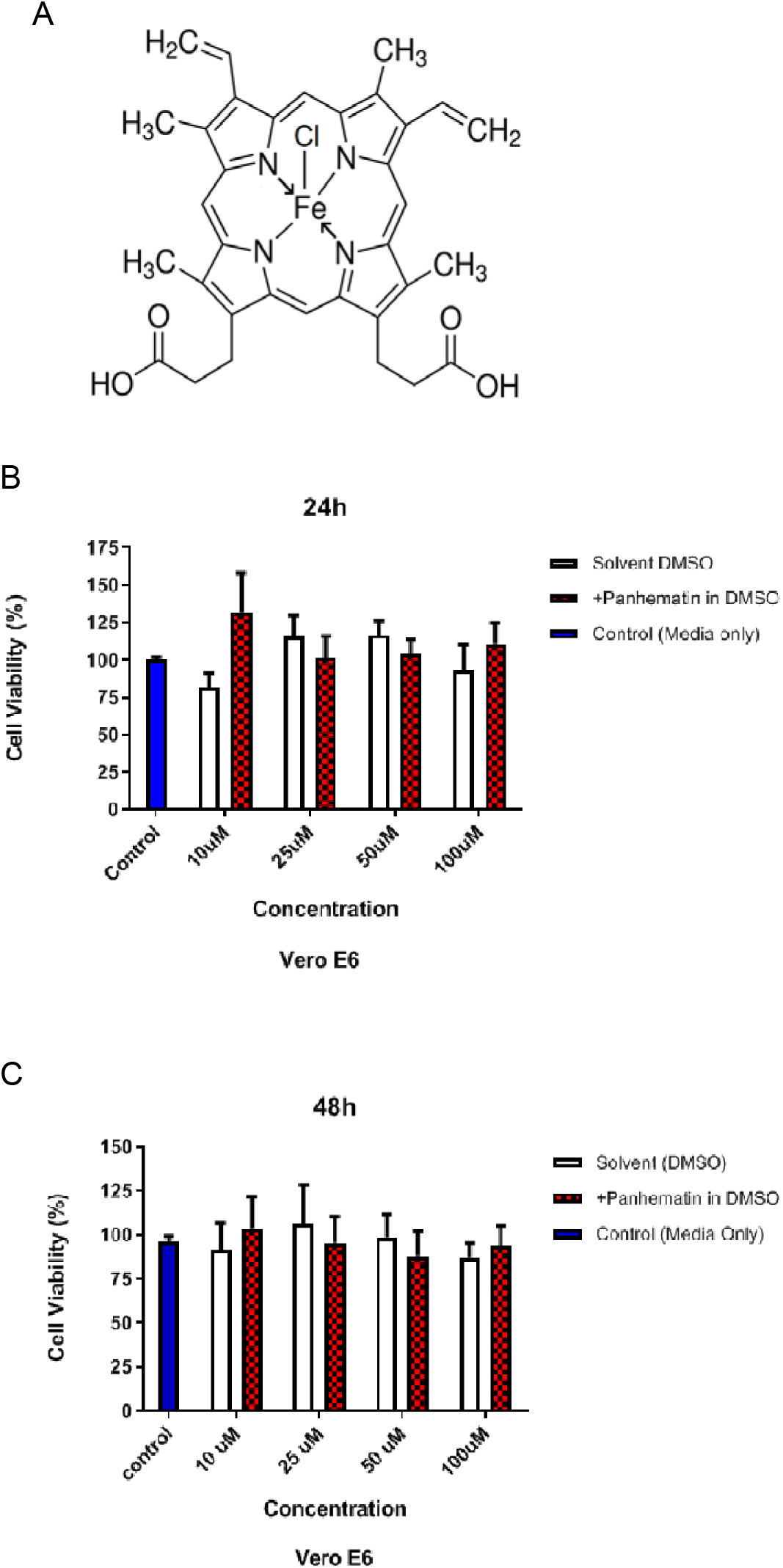
Hemin and its toxicity in Vero E6 cells. A) Strucure of Hemin. The viability of Vero E6 cells after the treatment with Hemin at the concentrations of 10, 25, 50, and 100 μM for A) 24 and B) 48 hrs. Data represent the mean and the SD of triplicate samples. Blue bars show the cells treated with the only cell culture media (Control). White bars show the cells treated with the cell culture media including DMSO as drug’s solvent (Solvent Control). Red bars show the cells treated with Hemin. * indicates p<0.05. All data are represented as the mean ± SD (n= 3)

The potential of HO-1 centric therapeutics to combat severe COVID-19 has been noted by several groups, yet the role of HO-1 in SARS-CoV-2 infection remains unclear (Dattilo, 2020; Fakhouri et al., 2020; Hooper, 2020; Maiti, 2020; Rossi et al., 2020; Singh et al., 2020; Wagener et al., 2020). Here we present experimental evidence for the antiviral capabilities of the HO-1-inducing drug Hemin in a cell line model of SARS-CoV-2 infection. We observed decreased infectivity in the presence of Hemin. We also observed a decrease in the expression and proteolytic activity of TMPRSS2, a critical mediator of cell entry by SARS-CoV-2, in cells treated with Hemin. These preliminary results, combined with expected physiological benefits of HO-1 induction, make Hemin a strong drug repurposing candidate for COVID-19 meriting further exploration *in vivo.*

## Materials and Methods

### Cell and Virus Culture

The cell culture media used for Vero E6 (African green monkey, kidney epithelial) cells consisted of high glucose Dulbecco’s modified Eagle medium (DMEM) (Capricorn) with 10 % fetal bovine serum (FBS) (Biolegend), 100 U/mL of penicillin (Lonza) and 0.1 mg/mL of streptomycin (Lonza). Cells were maintained in a humidified atmosphere at 37 °C and 5 % CO_2_. For the experiments control referred for non-treated cells, solvent control referred for cells treated with drug solvent DMSO.

Ank1, a local isolate of SARS-CoV-2, which was obtained from the biosafety level 3 (BSL3) laboratory based in the Department of Virology, Faculty of Veterinary Medicine, Ankara University. Viruses were propagated in Vero E6 cells using high glucose DMEM media containing 2% FBS and 1% penicillin and streptomycin.

### MTT Assay

MTT assay (Millipore, CT02) was used to analyze the toxic doses of Hemin in non-infected Vero E6 cell lines. Non-infected cells were plated on 96-well plates at a number of 10,000 cells/well and incubated 24h before treatment with Hemin. DMSO was used to dissolve Hemin and its serial dilutions (10, 25, 50, 100 μM) were prepared by diluting with DMEM. The cells were treated with the Hemin concentration gradient for 24 and 48 hours. Following 2 hours of incubation with MTT solution the absorption was measured at 570 nm using Epoch™Microplate Spectrophotometer to measure cell proliferation.

### Viral Plaque assay

A plaque assay was conducted using infected Vero E6 cells grown in 24-well plates. The cells were infected with 0.01 MOI of SARS-CoV-2 Ank1 isolate for 1 h at 37 <u>°</u>C. At the end of the incubation period, all virus suspension was removed from the cell surface and the cells were treated with high glucose (4g/L) Dulbecco’s Modified Essential Medium (DMEM) (2% Fetal Bovine Serum (BI, Israel) 1:100 antibiotic solution (BI, Israel)) and 2 or 20 μM Hemin. After 48h post-inoculation, cells were washed and fresh culture media was added onto the wells. After 96h of incubation culture mediums and cell layer were collected separately for infectivity quantification with a plaque assay. Experiments were performed in triplicates. At the end of infection, culture medium was collected and subjected to a freeze-thaw cycle. Thawed samples of infected culture medium and scraped Vero E6 cell layer were prepared in 10-fold dilutions in High Glucose DMEM without FBS a total of 200 μL of dilutions were inoculated on Vero E6 cells grown in a 24-well cell culture plate. The cells were incubated at 37<u>°</u>C for 1h to allow for attachment of viral particles. At the end of this period, cells were overlaid by 1.6% carboxymethyl cellulose (CMC-Sigma Aldrich, USA) and further incubated for 4 days under the same conditions. Infected and uninfected control cells were fixed with 3.5% formaldehyde (Sigma Aldrich, USA) for 20 minutes and stained with 0.75% Crystal Violet (Merck, USA). The number of plaques that occurred in each virus dilution was counted and the plaque-forming unit (PFU) was calculated as described previously (Hindawi et al., 2018). Results were evaluated statistically using the Two-way ANOVA (ordinary) technique (Prism 8 v8.2.1, GraphPad).

### Flow Cytometry

Flow cytometry (NovoCyte Flow Cytometry) assay was used to the TMPRSS2 protein expression in noninfected Vero E6 cells treated with Hemin. Vero E6 cells were harvested as control, solvent control and drug treatment groups with Trypsin-EDTA (2X) and single cell suspensions were stained using an antibody against TMPRSS2 (Santa Cruz, sc-515727). Samples were fixed with 4 % paraformaldehyde which was followed by permeabilization step or 15 min at room temperature. Data analysis was performed using NovoExpress 1.3.0 Software (ACEA Biosciences, Inc).

### mRNA expression by qRT-PCR

Quantitative real-time PCR (qRT-PCR) was used to determine the mRNA expression levels of TMPRSS2 and ACE2 in non-infected Vero E6 cells treated with Hemin. Total RNA was isolated with the Trizol (GeneAll® RiboEx™, 301-001). 1 μg of total RNA was used to synthesize cDNA (TONBO Biosciences, 31-5300-0100R). qRT-PCR analysis was carried out with TONBO Biosciences CYBERFast™ qPCRHi-ROX Master Mix. Gene expression was normalized to the GAPDH expression and assessed using the 2^-ΔΔCT^ method as fold difference compared to the solvent control.

### Western Blotting

Western blotting was used to obtain ACE2 protein expression in non-infected Vero E6 cells treated with Hemin. Protein lysates were prepared from the control, solvent control and 100μM Hemin treated cells collected with a scraper. The concentration of protein was calculated using BCA Protein Assay Reagent (Pierce, Rockford, IL) and 60 μg of each protein sample was loaded on the polyacrylamide gel and then transferred to polyvinylidenedifluoride (PVDF) membrane (TransBlot Turbo, Bio-Rad) with semi-dry transfer (Bio-Rad, Hercules, CA). The monoclonal anti-ACE2 antibody (ProSci, Poway, CA,1:750 diluted in 5% BSA blocking buffer) and the goat anti-rabbit IgG-horseradish peroxidase (HRP)-conjugated antibody (Santa Cruz Biotechnology, Dallas, TX, 1:2500) were used as primary and secondary antibodies, respectively, for ACE2 protein expression. The monoclonal anti-β-actin antibody (SantaCruz,1:1500) and the goat anti-mouse HRP conjugated (Pierce, Rockford, IL) were used as primary and secondary antibodies, respectively, for β-actin expression. Membrane images were developed with the ChemiDoc MP Imaging System (Bio-Rad, Hercules, CA).

### TMPRSS2 Proteolytic activity assay

The proteolytic activity of TMPRSS2 was evaluated in non-infected Vero E6 cells treated with Hemin. 20,000 cells/well were seeded into a 96-well plate in 100 μl of DMEM-HG-10%FBS-1%P/S. After 24 hours, cells were treated with Hemin (25, 50, 100 μM) for 48 hours. Cells were incubated with fluorogenic synthetic peptide Boc-Gln-Ala-Arg-AMC (Enzo, BML-P237-0005) at a final concentration of 200 μM at 37 °C for 30 minutes. Supernatants were separately collected from each well and centrifuged at low-speed for 10 min at 4 °C to remove the debris. Cleared supernatant was added to a new 96-well plate and, fluorescence intensity was measured using fluorescence spectrometers (Berthold Mithras LB, 943) at 380 nm and 460 nm.

### In silico modelling of drug-protein interactions

The X-ray structure of Hemin was downloaded from the CCDC (Cambridge Crystallographic Data Centre) (KOENIG, 1965) and an electrostatic potential (ESP) map was generated using the density functional theory (DFT) method at the B3LYP/6-31 G(d, p) level of theory using Gaussian 09W (Frisch et al. 2009) and the GausView program package (Dennington et al. 2009). The following PDB files were downloaded from the RCSB Protein Data Bank (PDB) (https://www.rcsb.org): 6VSB, 6VXX (open and closed forms of spike glycoproteins respectively), 6M03 (main protease), 6M71 (RNA polymerase), 5X29 (envelope protein), 6M0J (ACE2-bounded spike proteins), 1R42 (ACE2 protein) and 6EHA (Heme oxygenase 1). Before docking calculations, water and ligand molecules were removed from all protein structures using the BIOVIA Discovery Studio 2021 software (Dassault Systèmes 2021) *(BIOVIA Discovery Studio - BIOVIA - Dassault Systèmes®*, n.d.). For docking calculations, PDB files for proteins and Hemin were converted into PDBQT file format using default parameters of AutoDock Vina (ver.1.12) (Trott & Olson, 2009). The binding affinity energies (ΔG, kcal·mol^-1^) of Hemin with proteins were calculated with blind docking (supplementary material, DockingRawData.zip). The nine conformations with the highest affinity energy were taken into consideration. BIOVIA and UCSF Chimera ver. 1.15 (Pettersen et al., 2004) were used for two- and three-dimensional analysis and imaging of complexes. CAVITY (https://repharma.pku.edu.cn/cavity/home.jsp) (Yuan et al., 2013) was used to detect cavities and their druggability (supplementary material, DockingRawData.zip) in the prepared protein models.

### Statistical Analysis

MTT, flow cytometry, western blotting and proteolytic activity assays were performed in three biological replicates, and qRT-PCR assays were performed in six biological replicates. The paired two-tailed T test was used to compare the median values of the control and the drug-treated groups. Analysis of the variance was conducted on the replicate values of the experiment groups. P values <0.05 was accepted as statistically significant. * indicates that p<0.05. The data was analyzed using GraphPadPrism v 8.01.

## Results

### Hemin decreases SARS-CoV-2 infectivity in Vero E6 cells

No significant change was observed in cell viability with increasing concentration of Hemin (Figure 1B-C). The highest concentration (100 μM), at which there was no toxic effect on viability compared to the control, was selected for use in subsequent experiments. Following toxicity assessment, antiviral activity was assessed following SARS-CoV-2 infection *in vitro* in Vero E6 cells. Plaque assays showed a concentration dependent decrease of infectivity titers in the presence of Hemin, both in cell-free and cell-associated samples (Figure 2). This result suggests that Hemin shows antiviral activity against SARS-CoV-2.

**Figure 2:**
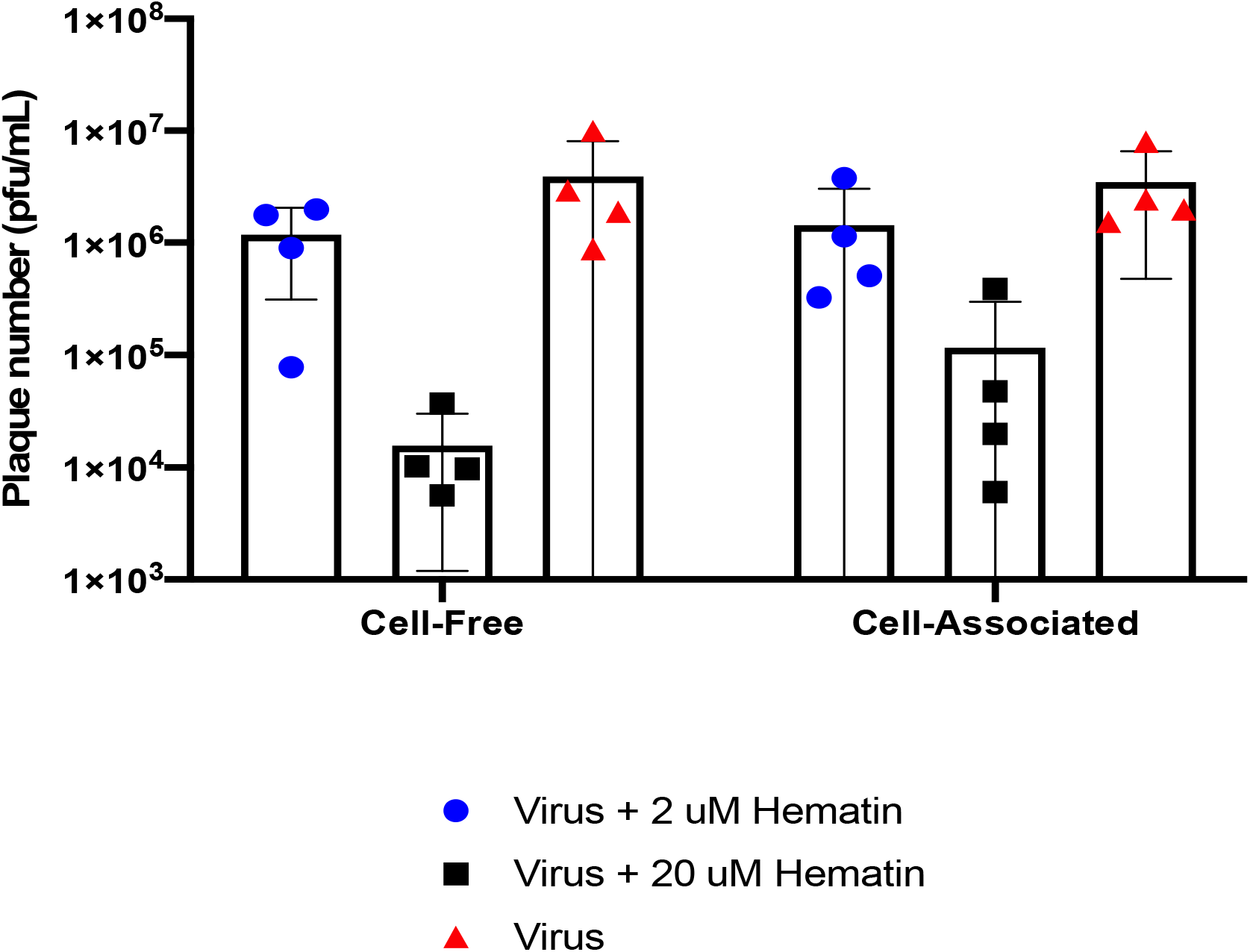
Antiviral activity of Hemin. Infectivity titers of SARS-CoV-2 virus cultured in the presence of 2 and 20 μM Hematin was plotted with individual dots and average bars. Data was obtained from A) cell-free B) cell-associated samples.

### Hemin decreases expression of viral entry mediators

qRT-PCR was used to evaluate whether Hemin affects the mRNA expression levels of TMPRSS2 and ACE2, which are critical mediators of virus entry (Bertram et al., 2013). Non-infected Vero E6 cells treated with 100 μM Hemin demonstrated a significant (~ 4-fold) decrease in expression of both TMPRSS2 and ACE2 mRNA expression after 48 hours (Figure 3). TMPRSS2 total protein in Vero E6 cells treated with Hemin (100 μM) was evaluated by flow cytometry. TMPRSS2 protein level was significantly decreased in the Hemin-treated group (8,23 ± 0,7%) compared to both control (87,20 ± 4,64 %) and solvent group (69,93 ± 1,25 %) (Figure 4A). ACE2 protein expression was evaluated with western blotting. There was no observable difference at protein levels in Vero E6 cells treated with Hemin (100 μM) for 48 hours compared to the control (Figure 4B). These results suggest that Hemin might interfere with virus entry as a consequence of diminished mRNA expression of TMPRSS2 and ACE2 or decreased total TMPRSS2 protein.

**Figure 3:**
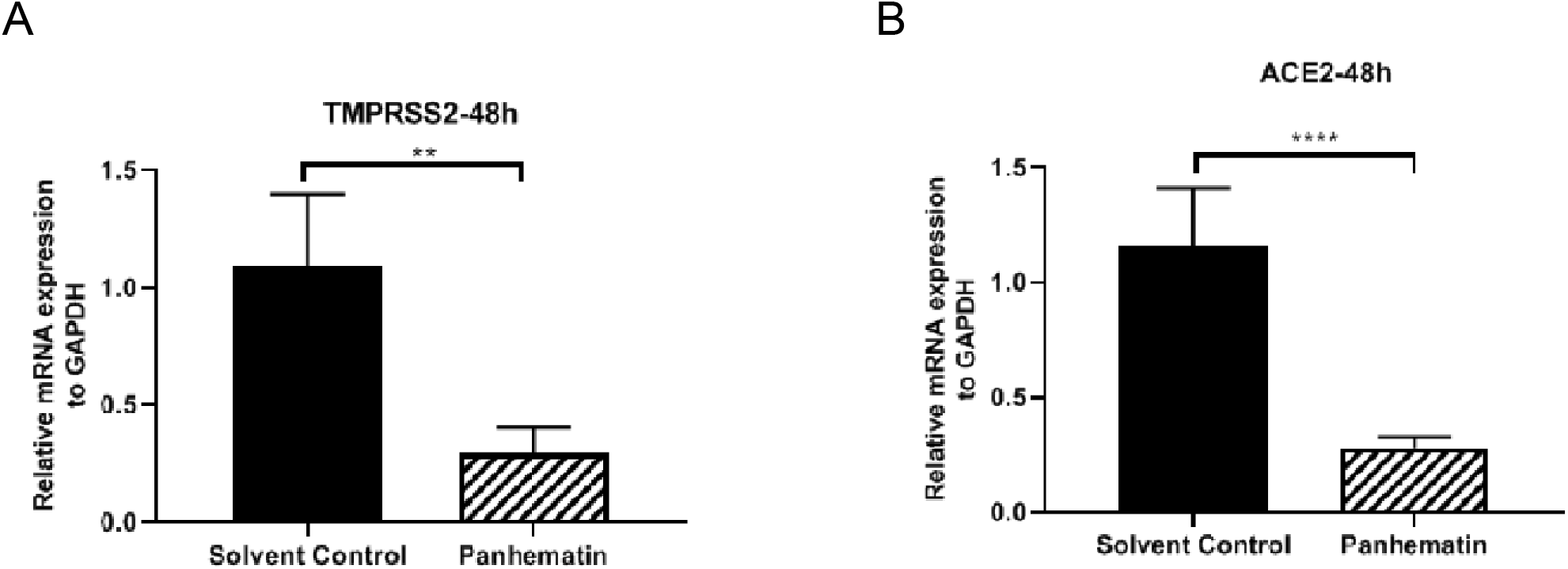
qPCR. TMPRSS2 and ACE2 mRNA expression levels in the absence and presence of the Hemin treatment of Vero E6 cells for 48 hours were demonstrated by qRT-PCR. (P<0.05). * indicates p<0.05. All data are represented as the mean ± SD (n= 6).

**Figure 4:**
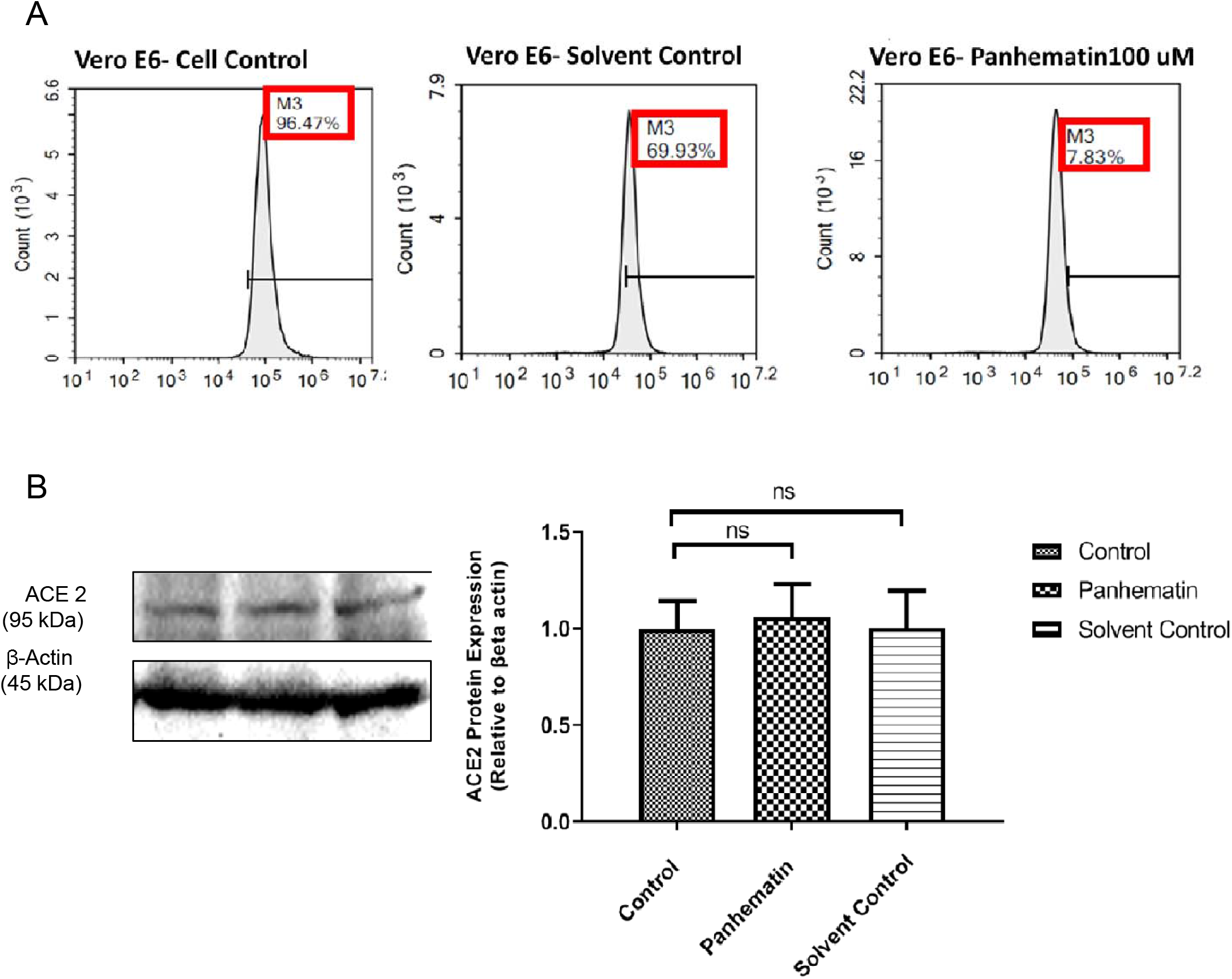
A) Flow Cytometry for TMPRSS2 protein expression. Histogram figures show the TMPRSS2 protein expression levels in the Vero E6 cells treated with 100 μM Hemin, solvent, and control for 48 hours. Hemin treated cells indicate a significant decrease in the TMPRSS2 expression (Paired Two-tailed T test, P<0.05). All data are represented as the mean ± SD (n= 3) **B) Western Blotting for ACE2 protein expression.** Hemin does not have any effect on the ACE2 protein expression level. The graph shows the analysis of band intensities indicating the protein levels of ACE2. Solvent treatment was used as a negative control. * indicates p<0.05. All data are represented as the mean ± SD (n= 3)

### Hemin suppresses the proteolytic activity of TMPRSS2

TMPRSS2 is a transmembrane serine protease that cleaves the spike protein of SARS-CoV-2 during membrane fusion between host cell and virus. Therefore, inhibition of TMPRSS2 protease activity was used as a proxy to measure the anti-viral effect of drugs at a molecular level (Shrimp et al.,2020; Sanderson et al., 2020). to examine the TMPRSS2-activated matriptase proteolytic activity The resonance energy transfer of a fluorescently-labeled synthetic protease substrate (Boc-Gln-Ala-Arg-AMC, ENZO Life Sciences) as described in Ko et al., 2015). Boc-Gln-Ala-Arg-AMC was added to Vero E6 cells treated with 25, 50, and 100 μM Hemin for 48h. A concentration gradient was used to determine the dose dependent effect of Hemin on the treated cells considering TMPRSS2 proteolytic activity. Hemin treatment exhibited significantly decreased TMPRSS2 proteolytic activity at concentrations of 50 and 100 μM compared to the control (Figure 5), suggesting that Hemin may diminish the proteolytic activity of TMPRSS2 in a dose dependent manner. This result could partially explain the underlying mechanism of anti-viral activity of Hemin at a molecular level.

**Figure 5:**
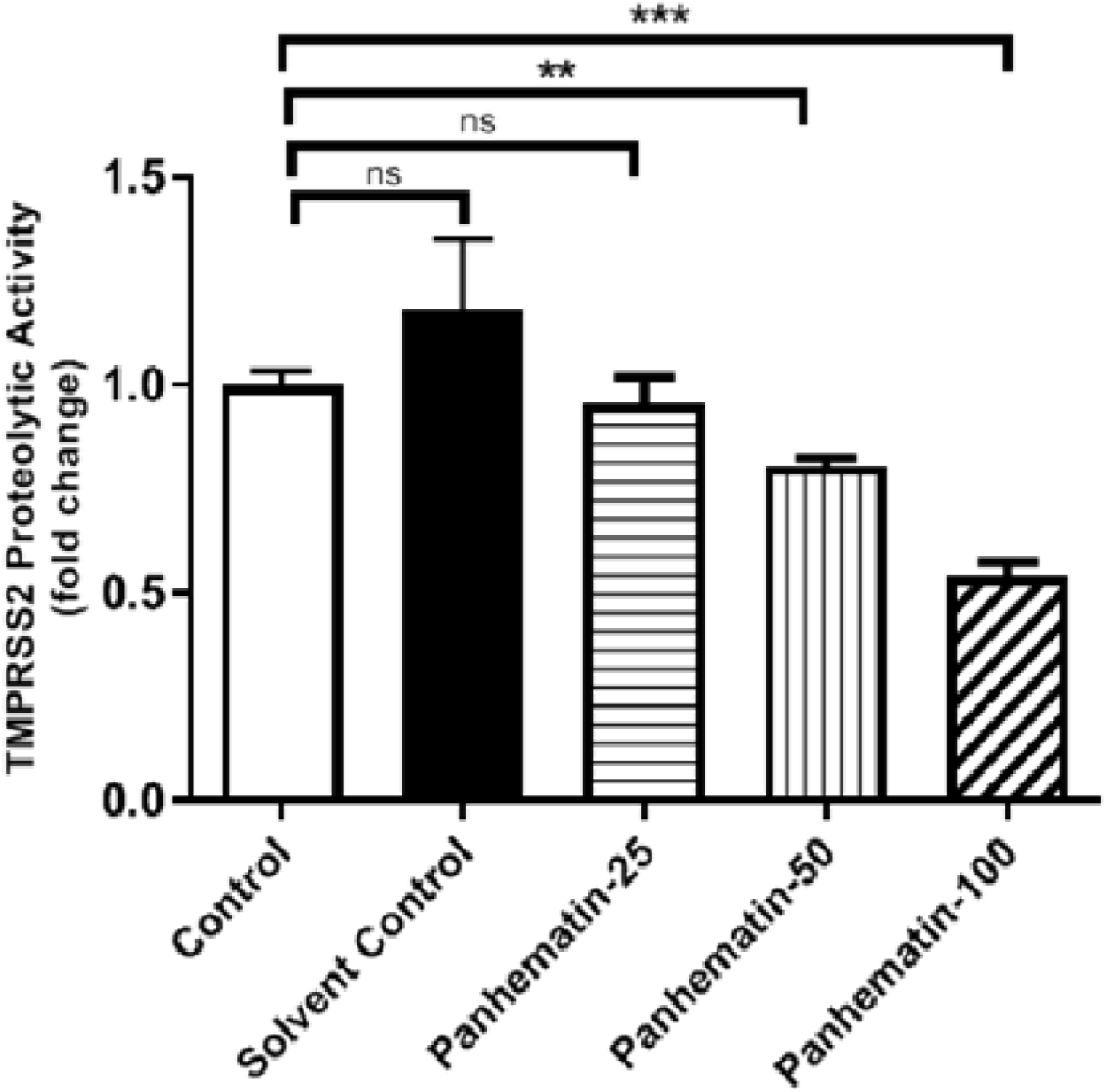
TMPRSS2 proteolytic activity. Examination of the proteolytic activity of TMPRSS2 enzyme was performed by the substrate assay analysis. The concentrations of Hemin were analysed compared to the control. * indicates p<0.05. ns indicates non-significance. All data are represented as the mean ± SD (n= 2).

### In silico analysis of Hemin with SARS-CoV-2 viral proteins

The ESP map of the hemin molecule shows the porphyrin ring and the acetate groups region as electrophilic and nucleophilic attack sites (Figure S1). This feature indicates that the molecule can act as a hydrogen donor and acceptor, interact non-bonding with amino acids and also can be held dynamically on the protein surface. When the affinity energies of hemin-protein complexes are examined (supplementary material, DockingRawData.zip); It is seen that the highest affinity value is - 9.2 kcal/mol against 6EHA (heme oxygenase 1) and the lowest affinity value with −5.5 kcal/mol against 5X29 (envelope) proteins (Figure S2a and Figure S2b respectively). The highest energy conformation of Hemin-6EHA complex is ΔG= −9.2 kcal/mol and the lowest energy conformation is ΔG= −7.3 kcal/mol. In the affected area, Hemin makes H-bond, van der Waals, Pi-Pi and Alkyl interactions with SER:142, LYS:179, GLY:139, HIS:25, PHE:207, etc. amino acids (Figure 6A). In addition, these interactions are identical to the amino acid interactions in the 6EHA-Hemin complex which given in the PDB database. This shows the accuracy of the calculated 6EHA-Hemin complex model. When the chemical environment of hemin on the protein surface is analyzed, Figure 6B shows its behavior as a hydrogen donor and acceptor, Figure 6C the hydrophobicity, and Figure 6D the charge distribution of its environment. One of the acetate group of hemin forms van der Waals bonds with amino acids in the hydrophobic region which (GLN: 145, VAL: 146, LEU: 147) indicates that it can hold on to the protein surface. On the other hand, considering role of the 5X29 envelope protein (https://www.uniprot.org/uniprot/P59637), hemin’s moderate affinity for the 5X29 envelope protein - 6.0 kcal/mol is also important. The hydrogen bond, van der Waals interactions and other non-covalent interactions of hemin, especially with amino acids located in the hydrophobic region of the 5X29 protein (Figure S3), show the effect of hemin on the envelope protein. According to UniProtKB database (https://www.uniprot.org/uniprot/P0DTC2), 6VYB (open form) and 6VXX (close form) spike proteins have two binding areas. These are amino acids between 319-541 (Receptor Binding Domain, RBD) and 437-508 (receptor-binding motif). Besides, 30-41, 82-84 and 353-357 amino acids are active in binding of ACE2 protein (1R42) to spike proteins https://www.uniprot.org/uniprot/Q9BYF1). When the picture shots are examined (Figure S4A and S4B), it is seen that Hemin shows conventional hydrogen bonds (as donor and acceptor), van der Waals interactions and other non-covalent interactions with protein amino acids. The highest druggability score of active region of the 6VSB and 6VXX Spike proteins corresponds to the middle region of the proteins. The highest affinity conformations of Hemin are also located in these regions (Figure S4Aa, S4Ab). However, the 6VSB protein has two conformations (mode 4 and 8) in the RBD (Receptor Binding Domain) region with affinity energies of −8.0 kcal/mol and −7.6 kcal/mol, respectively. The 3D and 2D representation of the interaction of these conformations are given in Figure S4Be-k. The hemin 6M0J (Spike-ACE2 bounded protein) has an affinity both the ACE2 and Spike-ACE2 interface between −8.0 kcal/mol and −7.1 kcal/mol (Figure S4Bd). However, the affinity of the 4^th^, 6^th^, 7^th^, and 9^th^ conformations of hemin to the interface of the 6M0J Spike-ACE2 bounded protein is also shown in Figure S4B1. When the 2D and 3D representations of the conformer (Figure S4Bm-x) are analyzed, it is seen that it makes non-covalent interactions with amino acids that are important in Spike-ACE2 interaction. It can also be said that the charge distribution in the impact region is neutral and it interacts in the area where solvent accessibility is low (Figure S4q-t). In the ACE2 (1R42)-hemin complex, all the calculated conformations of hemin are located in the druggable cavity region of the protein (Figure S4Ac). It is also important that the third conformation of hemin with an energy of −7.6 kcal/mol interacts with the amino acid PHE:40, which is among the amino acids interacting with Spike (Figure S4Bk).

**Figure 6.**
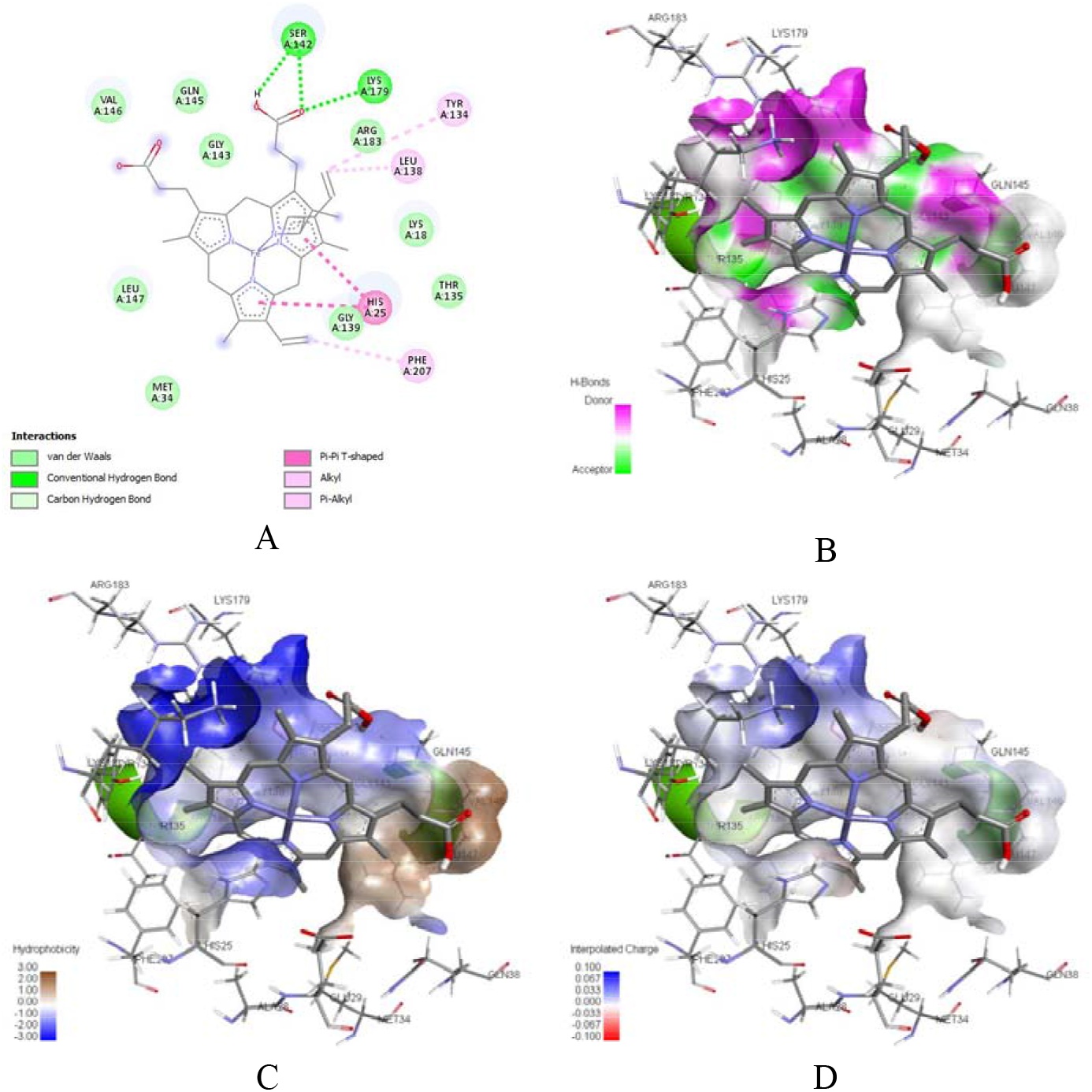
In silico docking analysis of Hemin against major viral proteins. The highest energy conformation of the Hemin-6EHA complex A) 3D interaction, B) hydrogen donor-acceptor, C) hydrophobicity and D) interpolated charge properties

6M03 and 6M71 are main proteases and RNA polymerase proteins, respectively. Hemin’s affinity for 6M03 protein varies between −8.0 kcal/mol and −7.4 kcal/mol (Figure S5a). The highest-energy conformation of Hemin is located in the region where 6M03 with the highest druggability score. When Figure S5b and S5e are examined together, hemin give hydrogen bond with GLU:288, which is located in the hydrophobic region of the protein, and also with other amino acids; LEU:271, LEU:272, GLY:275, MET:276, LEU:286, LEU:287 give van der Waals interactions. This is related to the persistence of the hemin protein on the surface. Indeed, the SAS map of the 6M03 protein (Figure S5d) shows that hemin can give van der Waals interactions and Pi-Alkyl interactions with amino acids ARG:131, TRY:239, LEU:271 and ASP:289 in the lowest SAS region of the protein. This positively affects the interaction of hemin with protein and increases the residence time of hemin on the protein surface. Figure S5c shows that heme can act as both a hydrogen donor and acceptor on the protein surface. The results of the interaction of hemin with the 6M71 protein are also given in Figure S6. The effect and properties of the highest conformation of hemin are similar to that of other proteins. On the other hand, unlike other proteins, hemin shows affinity for almost every region of the 6M71 protein between −7.8 kcal/mol and −7.2 kcal/mol (Figure S6a).

## Discussion

Commercially available Hemin_®_ (Ovation Pharmaceuticals, Inc., USA) has been used in clinic for acute Intermittent porphyria (AIP) since 1983. Hemin is a heme analog that is well known to activate the heme catabolizing enzyme HO-1 both *in vitro* and *in vivo* (X. Chen et al., 2018). HO-1 and heme metabolites have strong antiviral properties (Singh et al., 2020).They also protect against damaging inflammation and coagulopathy (Nath et al., 2020; B. Wu et al., 2019) Dysregulated inflammation and coagulopathy are particularly insidious features of COVID-19 that cause widespread damage to multiple organ systems (Tay et al., 2020). Many have suggested that induction of the HO-1 system by various agents could be of great benefit to patients suffering from COVID-19, (Dattilo, 2020; Fakhouri et al., 2020; Hooper, 2020; Maiti, 2020; Rossi et al., 2020; Singh et al., 2020; Wagener et al., 2020) however this idea is yet to be validated by experimental evidence. Relevant models in which Hemin has been demonstrated as protective include; acute lung injury (X. Chen et al., 2018; Chi et al., 2016; Luo et al., 2014), inflammation (Kim & Lee, 2013; Konrad et al., 2016; Vitali et al., 2020; Yamauchi et al., 2004), influenza pneumonia (Wang et al., 2017), ischemia reperfusion injury (Attuwaybi et al., 2004; Lakkisto et al., 2009; Rossi et al., 2018; Xue et al., 2007), acute hepatic injury (Wen et al., 2007), vascular thrombosis (Desbuards et al., 2007; Nath et al., 2020), and septic muscle wasting (Yu et al., 2018). Preclinical *in vitro* and *in vivo* studies are needed to demonstrate the effect of HO-1 inducers on SARS-CoV-2 infection and to better understand the possible therapeutic utility of these drugs. In the present study, we captured the antiviral effect of Hemin *in vitro* to evaluate the drug as a possible therapeutic strategy against COVID-19.

The antiviral potential of hemin against SAR-CoV-2 was demonstrated by a dose dependent decrease in infectivity of the virus in Vero E6 cells. To determine whether the observed decrease in infectivity could be due to hindrance of viral entry, we examined the expression profiles of two critical mediators of viral entry, namely TMPRSS2 and ACE2 (McKee et al., 2020). In Vero E6 cells treated with Hemin, we observed a significant decrease in mRNA expression of both TMPRSS2 and ACE2 relative to cells not treated with Hemin (Figure 3). We also observed a significant decrease in TMPRSS2 total protein (Figure 4A) and protease activity (Figure 5) in hemin treated cells relative to controls. Interestingly, ACE2 total protein was consistent between Hemin treated and control cells (Figure 4B). Taken together, these results support the possibility that antagonism of viral entry mediators, particularly TMPRSS2, was responsible for the observed decrease in infectivity. The HO-1 system exhibits antiviral activity through diverse mechanisms in a broad range of viral diseases (Devadas & Dhawan, 2006; Espinoza et al., 2017; Lu et al., 2020; Protzer et al., 2007; Tseng et al., 2016; Wang et al., 2017). To our knowledge, suppression of viral entry by TMPRSS2 antagonism has yet to be reported as an antiviral mechanism of the HO-1 system. The link between HO-1 and TMPRSS2 remains incompletely resolved, but there is a possible explanation through the androgen receptor (AR) pathway. Expression of TMPRSS2 is regulated with Androgen Receptor (AR) pathway. AR pathway is activated by testosterone and dihydrotestosterone molecules (Feng & He, 2019). Activation of AR pathway induces translocation of AR dependent transcription factor to nucleus and induces TMPRSS2 expression (Tan et al., 2015). HO-1 induction decreases the activity of AR leading to a decrease in the expression of TMPRSS2 (Elguero et al., 2012). In addition, the expression of TMPRSS2 increases with the presence of ROS whereas induction of HO-1 via Hemin inhibits the production of ROS (Paszti-Gere et al., 2015). Our experimentalresults showed the dramatic decrease in both activity and the expression of TMPRSS2. Therefore, these results suggest that the expression of TMPRSS2 can be downregulated due to the decreased AR pathway activity and ROS production induced by HO-1 with Hemin. Future studies will be needed to elucidate the link between Hemin and TMPRSS2.

Clinical measurements of heme and HO-1 in COVID-19 patients indicate that dysregulation of heme catabolism is a feature of COVID-19. Dysfunctional heme catabolism is common in the critically ill, particularly in those with sepsis (Larsen et al., 2010). A small cohort study indicated elevation of heme and HO-1 levels in COVID-19 patients exhibiting oxygen desaturation relative to COVID-19 patients who were not oxygen desaturated (Su et al., 2021). Another study reported abnormal levels of porphyrins in the serum of COVID-19 patients (San Juan et al., 2020). Finally, the network of scavengers of heme, iron, and hemoproteins are also affected by COVID-19 infection (Yağcı et al., 2021). It is unclear whether heme catabolism is uniquely affected by COVID-19, however a novel interaction between HO-1 and the open reading frame 3a (ORF3a) of SARS-CoV-2 has been reported (Gordon et al., 2020). The interaction has not been mechanistically characterized however it is certainly plausible that SARS-CoV-2 has evolved to suppress the antiviral activities of HO-1. Accumulating evidence for dysregulation of heme catabolism and related pathways underscore the importance of interrogating therapeutics like Hemin which target these systems.

One of the major protective mechanisms of Hemin-mediated HO-1 induction is the attenuation of the NLRP3 inflammasome (Figure 7). Inflammasomes are activated by injury, infection, and stress, and they engage innate immunity through the activation of inflammatory caspases which drive the maturation and secretion of proinflammatory cytokines (Farag et al., 2020; C. Wu et al., 2019). The NLRP3 inflammasome is thought to be a major driver of inflammation, tissue damage, and coagulopathy in COVID-19 (Freeman & Swartz, 2020; Ratajczak & Kucia, 2020; van den Berg & te Velde, 2020). Furthermore, the close relatives of SARS-CoV-2, SARS-CoV and MERS-CoV, directly provoke the NLRP3 inflammasome (Castaño-Rodriguez et al., 2018; I. Y. Chen et al., 2019; Nieto-Torres et al., 2015; Siu et al., 2019). In macrophages, Hemin prevents inflammasome assembly by enhancing autophagy of ASC, a critical component of the inflammasome (Nurmi et al., 2017). Hemin treatment also decreases the expression of NLRP3 inflammasome components and by extension the proinflammatory cytokines IL-1B and IL-18 (Kim & Lee, 2013; Luo et al., 2014).

**Figure 7:**
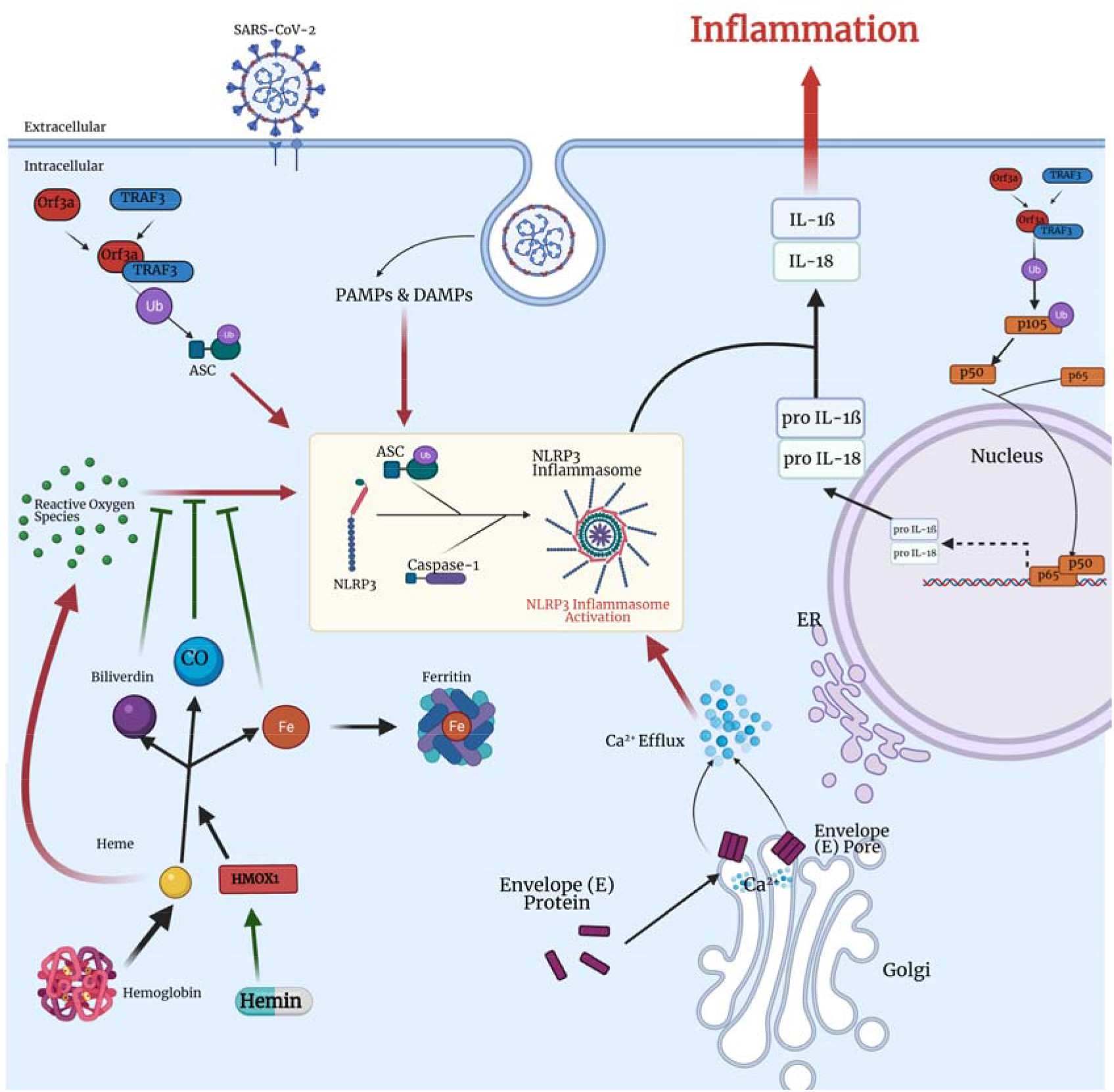
Antioxidant HMOX1 pathway and NLRP3 inflammasome assembly induced by SARS-CoV-2 enntry. Inflammasomes are part of innate immunity which are activated due to viral/bacterial infection or stress. Activation of inflammasomes engage innate immune response including activation of inflammatory caspases and secretion of proinflammatory cytokines. Nod-like receptor family, pyrin domain-containing 3 (NLRP3) inflammasome complex is triggered by presence of reactive oxygen species (ROS) and activation of pattern recognition receptors (PRRs) - Toll like receptors (TLRs)-with recognition of damage-associated molecular patterns (DAMPs) and pathogen-associated molecular patterns (PAMPs). NLRP3 inflammasome assembly starts with association of NLRP3 protein to adaptor ASC protein and caspase-1 protein leading oligomerization of NLRP3 inflammasome (Swanson et al., 2019). Caspase-1 protein in the complex cleaves pro-IL-1β and pro-IL-18 into IL-1β and IL-18 respectively to induce inflammation. Orf8a, Orf3a, and Envelope (E) proteins of SARS-CoV-2 induces the activation of NLRP3 inflammasome (I. Y. Chen et al., 2019; Nieto-Torres et al., 2015). TNF receptor-associated factor 3 (TRAF3) interaction with Orf3a protein induces ubiquitination of ASC adaptor protein for the inflammasome assembly and activates NF-κB signaling pathway to increase the expression of pro-IL-1β and pro-IL-18 (Siu et al., 2019). Higher levels of IL-1β and IL-18 indicated in patients’ blood, lungs and lymphoid tissues for SARS-CoV-2 studies suggesting activation of NLRP3 inflammasome complex (Freeman & Swartz, 2020). The activity of NLRP3 inflammasome can be downregulated by inducing antioxidant activity of HMOX1 pathway with FDA approved drug Hemin (Hemin) (Luo et al., 2014). Hemin decreases the expression of ASC, NLRP3 and caspase-1 proteins by inducing the activity of HO-1 leading inhibition of NLRP3 inflammasome and reduction of oxidative stress, inflammatory cytokines - TNF-α, IL-1β, IL-18 (Yoshihara et al., 2013). Also, Hemin interfere with inflammasome assembly with prevention of NLRP3, ASC and caspase-1 association (Kim & Lee, 2013). Created with BioRender.com.

The complexity of living organisms makes it difficult for *in vitro* systems to fully capture the effect of drug perturbation. Much of the benefit of the HO-1 system relies on its ability to modulate the immune response and mitigate oxidative damage to host tissues, which could not be captured by our preliminary experiments (Gozzelino et al., 2010). However, we expect that the anti-inflammatory and antiviral benefits of HO-1 induction will be further exemplified by future studies *in vivo.* Models in which immune system dynamics and physiological changes can be observed will likely be necessary fully capture the effect of Hemin on SARS-CoV-2 infection. Importantly, the timing of Hemin administration may be of critical importance to the efficacy of the drug, which will be particularly important to account for in future experiments (Rossi et al., 2018).

## Supporting information

Supplemental Raw Data

## Acknowledgement

The research for this paper was financially supported by the Scientific and Technological Research Council of Turkey (TUBITAK) under grant number 18AG020. We thank TURKSAT for allowing us to use their server infrastructure and the Ankara University Information Technologies Department for technical support.

## References

Attuwaybi, B. O., Kozar, R. A., Moore-Olufemi, S. D., Sato, N., Hassoun, H. T., Weisbrodt, N. W., & Moore, F. A. (2004). Heme oxygenase-1 induction by hemin protects against gut ischemia/reperfusion injury. Journal of Surgical Research, 118(1), 53–57. https://doi.org/10.1016/j.jss.2004.01.010

BIOVIA Discovery Studio - BIOVIA - Dassault Systèmes®. (n.d.). Retrieved May 9, 2021, from https://www.3ds.com/products-services/biovia/products/molecular-modeling-simulation/biovia-discovery-studio/

Castaño-Rodriguez, C., Honrubia, J. M., Gutiérrez-Álvarez, J., DeDiego, M. L., Nieto-Torres, J. L., Jimenez-Guardeño, J. M., Regla-Nava, J. A., Fernandez-Delgado, R., Verdia-Báguena, C., Queralt-Martín, M., Kochan, G., Perlman, S., Aguilella, V. M., Sola, I., & Enjuanes, L. (2018). Role of severe acute respiratory syndrome coronavirus viroporins E, 3a, and 8a in replication and pathogenesis. MBio, 9(3). https://doi.org/10.1128/mBio.02325-17

Chen, I. Y., Moriyama, M., Chang, M. F., & Ichinohe, T. (2019). Severe acute respiratory syndrome coronavirus viroporin 3a activates the NLRP3 inflammasome. Frontiers in Microbiology, 10(JAN), 50. https://doi.org/10.3389/fmicb.2019.00050

Chen, X., Wang, Y., Xie, X., Chen, H., Zhu, Q., Ge, Z., Wei, H., Deng, J., Xia, Z., & Lian, Q. (2018). Heme oxygenase-1 reduces sepsis-induced endoplasmic reticulum stress and acute lung injury. Mediators of Inflammation, 2018. https://doi.org/10.1155/2018/9413876

Chi, X., Guo, N., Yao, W., Jin, Y., Gao, W., Cai, J., & Hei, Z. (2016). Induction of heme oxygenase-1 by hemin protects lung against orthotopic autologous liver transplantation-induced acute lung injury in rats. Journal of Translational Medicine, 14(1), 35. https://doi.org/10.1186/s12967-016-0793-0

Dennington, R., Keith, T., and Millam, J. 2009. Gaussian 09W and Gauss-View, version 5. Semi-chem Inc., Shawnee Mission, Kansas, USA. Availablefrom http://gaussian.com.

Dattilo, M. (2020). The role of host defences in Covid 19 and treatments thereof. In Molecular Medicine (Vol. 26, Issue 1, pp. 1–15). BioMed Central Ltd. https://doi.org/10.1186/s10020-020-00216-9

Desbuards, N., Rochefort, G. Y., Schlecht, D., Machet, M.-C., Halimi, J.-M., Eder, V., Hyvelin, J.-M., & Antier, D. (2007). Heme oxygenase-1 inducer hemin prevents vascular thrombosis. Thrombosis and Haemostasis, 98(3), 614–620.

Devadas, K., & Dhawan, S. (2006). Hemin Activation Ameliorates HIV-1 Infection via Heme Oxygenase-1 Induction. The Journal of Immunology, 176(7), 4252–4257. https://doi.org/10.4049/jimmunol.176.7.4252

Elguero, B., Gueron, G., Giudice, J., Toscani, M. A., de Luca, P., Zalazar, F., Coluccio-Leskow, F., Meiss, R., Navone, N., de Siervi, A., & Vazquez, E. (2012). Unveiling the association of STAT3 and HO-1 in prostate cancer: Role beyond heme degradation. Neoplasia (United States), 14(11), 1043–1056. https://doi.org/10.1593/neo.121358

Espinoza, J. A., León, M. A., Céspedes, P. F., Gómez, R. S., Canedo-Marroquín, G., Riquelme, S. A., Salazar-Echegarai, F. J., Blancou, P., Simon, T., Anegon, I., Lay, M. K., González, P. A., Riedel, C. A., Bueno, S. M., & Kalergis, A. M. (2017). Heme Oxygenase-1 Modulates Human Respiratory Syncytial Virus Replication and Lung Pathogenesis during Infection. The Journal of Immunology, 199(1), 212–223. https://doi.org/10.4049/jimmunol.1601414

Fakhouri, E. W., Peterson, S. J., Kothari, J., Alex, R., Shapiro, J. I., & Abraham, N. G. (2020). Genetic polymorphisms complicate covid-19 therapy: Pivotal role of ho-1 in cytokine storm. In Antioxidants (Vol. 9, Issue 7, pp. 1–23). MDPI AG. https://doi.org/10.3390/antiox9070636

Farag, N. S., Breitinger, U., Breitinger, H. G., & El Azizi, M. A. (2020). Viroporins and inflammasomes: A key to understand virus-induced inflammation. In International Journal of Biochemistry and Cell Biology (Vol. 122, p. 105738). Elsevier Ltd. https://doi.org/10.1016/j.biocel.2020.105738

Feng, Q., & He, B. (2019). Androgen Receptor Signaling in the Development of Castration-Resistant Prostate Cancer. In Frontiers in Oncology (Vol. 9, p. 858). Frontiers Media S.A. https://doi.org/10.3389/fonc.2019.00858

Freeman, T. L., & Swartz, T. H. (2020). Targeting the NLRP3 Inflammasome in Severe COVID-19. In Frontiers in Immunology (Vol. 11, p. 1518). Frontiers Media S.A. https://doi.org/10.3389/fimmu.2020.01518

Frisch, M.J., Trucks, G.W., Schlegel, H.B., Scuseria, G.E., Robb, M.A., Cheeseman, J.R., Scalmani, G., Barone, V., Mennucci, B., Petersson, G.A., et al. 2009. Gaussian G09. Gaussian Inc., Wallingford, CT, USA. Available from https://gaussian.com/glossary/g09/.

Gordon, D. E., Jang, G. M., Bouhaddou, M., Xu, J., Obernier, K., White, K. M., O’Meara, M. J., Rezelj, V. V., Guo, J. Z., Swaney, D. L., Tummino, T. A., Hüttenhain, R., Kaake, R. M., Richards, A. L., Tutuncuoglu, B., Foussard, H., Batra, J., Haas, K., Modak, M., … Krogan, N. J. (2020). A SARS-CoV-2 protein interaction map reveals targets for drug repurposing. Nature, 583(7816), 459–468. https://doi.org/10.1038/s41586-020-2286-9

Gozzelino, R., Jeney, V., & Soares, M. P. (2010). Mechanisms of cell protection by heme Oxygenase-1. In Annual Review of Pharmacology and Toxicology (Vol. 50, pp. 323–354). Annual Reviews. https://doi.org/10.1146/annurev.pharmtox.010909.105600

Hindawi, S. I., Hashem, A. M., Damanhouri, G. A., El-Kafrawy, S. A., Tolah, A. M., Hassan, A. M., & Azhar, E. I. (2018). Inactivation of Middle East respiratory syndrome-coronavirus in human plasma using amotosalen and ultraviolet A light. Transfusion, 58(1), 52–59. https://doi.org/10.1111/trf.14422

Hooper, P. L. (2020). COVID-19 and heme oxygenase: novel insight into the disease and potential therapies. Cell Stress and Chaperones, 25(5), 707–710. https://doi.org/10.1007/s12192-020-01126-9

Kim, S. J., & Lee, S. M. (2013). NLRP3 inflammasome activation in d-galactosamine and lipopolysaccharide-induced acute liver failure: Role of heme oxygenase-1. Free Radical Biology and Medicine, 65, 997–1004. https://doi.org/10.1016/j.freeradbiomed.2013.08.178

Ko, C. J., Huang, C. C., Lin, H. Y., Juan, C. P., Lan, S. W., Shyu, H. Y., Wu, S. R., Hsiao, P. W., Huang, H. P., Shun, C. T., & Lee, M. S. (2015). Androgen-induced TMPRSS2 activates matriptase and promotes extracellular matrix degradation, prostate cancer cell invasion, tumor growth, and metastasis. Cancer Research, 75(14), 2949–2960. https://doi.org/10.1158/0008-5472.CAN-14-3297

Koenig, D. F. (1965). THE STRUCTURE OF ALPHA-CHLOROHEMIN. Acta Crystallographica, 18(4), 663–673. https://doi.org/10.1107/S0365110X65001536

Konrad, F. M., Knausberg, U., Höne, R., Ngamsri, K. C., & Reutershan, J. (2016). Tissue heme oxygenase-1 exerts anti-inflammatory effects on LPS-induced pulmonary inflammation. Mucosal Immunology, 9(1), 98–111. https://doi.org/10.1038/mi.2015.39

Lakkisto, P., Csonka, C., Fodor, G., Bencsik, P., Voipio-Pulkki, L. M., Ferdinandy, P., & Pulkki, K. (2009). The heme oxygenase inducer hemin protects against cardiac dysfunction and ventricular fibrillation in ischaemic/reperfused rat hearts: Role of connexin 43. Scandinavian Journal of Clinical and Laboratory Investigation, 69(2), 209–218. https://doi.org/10.1080/00365510802474392

Larsen, R., Gozzelino, R., Jeney, V., Tokaji, L., Bozza, F. A., Japiassú, A. M., Bonaparte, D., Cavalcante, M. M., Chora, Â., Ferreira, A., Marguti, I., Cardoso, S., Sepúlveda, N., Smith, A., & Soares, M. P. (2010). A central role for free heme in the pathogenesis of severe sepsis. Science Translational Medicine, 2(51), 51ra71–51ra71. https://doi.org/10.1126/scitranslmed.3001118

Lehmann, E., El-Tantawy, W. H., Ocker, M., Bartenschlager, R., Lohmann, V., Hashemolhosseini, S., Tiegs, G., & Sass, G. (2010). The heme oxygenase 1 product biliverdin interferes with hepatitis C virus replication by increasing antiviral interferon response. Hepatology, 51(2), 398–404. https://doi.org/10.1002/hep.23339

Lu, W., Shi, L., Gao, J., Zhu, H., Hua, Y., Cai, J., Wu, X., Wan, C., Zhao, W., & Zhang, B. (2020). Piperlongumine Inhibits Zika Virus Replication In vitro and Promotes Up-Regulation of HO-1 Expression, Suggesting An Implication of Oxidative Stress. Virologica Sinica, 1–11. https://doi.org/10.1007/s12250-020-00310-6

Luo, Y. P., Jiang, L., Kang, K., Fei, D. S., Meng, X. L., Nan, C. C., Pan, S. H., Zhao, M. R., & Zhao, M. Y. (2014). Hemin inhibits NLRP3 inflammasome activation in sepsis-induced acute lung injury, involving heme oxygenase-1. International Immunopharmacology, 20(1), 24–32. https://doi.org/10.1016/j.intimp.2014.02.017

Maiti, B. K. (2020). Heme/Hemeoxygenase-1 System Is a Potential Therapeutic Intervention for COVID-19 Patients with Severe Complications. In ACS Pharmacology and Translational Science (Vol. 3, Issue 5, pp. 1032–1034). American Chemical Society. https://doi.org/10.1021/acsptsci.0c00136

McKee, D. L., Sternberg, A., Stange, U., Laufer, S., & Naujokat, C. (2020). Candidate drugs against SARS-CoV-2 and COVID-19. In Pharmacological Research (Vol. 157, p. 104859). Academic Press. https://doi.org/10.1016/j.phrs.2020.104859

Nagasawa, R., Hara, Y., Murohashi, K., Aoki, A., Kobayashi, N., Takagi, S., Hashimoto, S., Kawana, A., & Kaneko, T. (2020). Serum heme oxygenase-1 measurement is useful for evaluating disease activity and outcomes in patients with acute respiratory distress syndrome and acute exacerbation of interstitial lung disease. BMC Pulmonary Medicine, 20(1). https://doi.org/10.1186/s12890-020-01341-1

Nath, K. A., Grande, J. P., Belcher, J. D., Garovic, V. D., Croatt, A. J., Hillestad, M. L., Barry, M. A., Nath, M. C., Regan, R. F., & Vercellotti, G. M. (2020). Antithrombotic effects of heme-degrading and heme-binding proteins. American Journal of Physiology. Heart and Circulatory Physiology, 318(3), H671–H681. https://doi.org/10.1152/ajpheart.00280.2019

Nieto-Torres, J. L., Verdiá-Báguena, C., Jimenez-Guardeño, J. M., Regla-Nava, J. A., Castaño-Rodriguez, C., Fernandez-Delgado, R., Torres, J., Aguilella, V. M., & Enjuanes, L. (2015). Severe acute respiratory syndrome coronavirus E protein transports calcium ions and activates the NLRP3 inflammasome. Virology, 485, 330–339. https://doi.org/10.1016/j.virol.2015.08.010

Nurmi, K., Kareinen, I., Virkanen, J., Rajamäki, K., Kouri, V.-P., Vaali, K., Levonen, A.-L., Fyhrquist, N., Matikainen, S., Kovanen, P. T., & Eklund, K. K. (2017). Hemin and Cobalt Protoporphyrin Inhibit NLRP3 Inflammasome Activation by Enhancing Autophagy: A Novel Mechanism of Inflammasome Regulation. Journal of Innate Immunity, 9(1), 65–82. https://doi.org/10.1159/000448894

Olagnier, D., Peri, S., Steel, C., van Montfoort, N., Chiang, C., Beljanski, V., Slifker, M., He, Z., Nichols, C. N., Lin, R., Balachandran, S., & Hiscott, J. (2014). Cellular Oxidative Stress Response Controls the Antiviral and Apoptotic Programs in Dengue Virus-Infected Dendritic Cells. PLoS Pathogens, 10(12), 1004566. https://doi.org/10.1371/journal.ppat.1004566

Paszti-Gere, E., Barna, R. F., Kovago, C., Szauder, I., Ujhelyi, G., Jakab, C., Meggyesházi, N., & Szekacs, A. (2015). Changes in the Distribution of Type II Transmembrane Serine Protease, TMPRSS2 and in Paracellular Permeability in IPEC-J2 Cells Exposed to Oxidative Stress. Inflammation, 38(2), 775–783. https://doi.org/10.1007/s10753-014-9988-9

Pettersen, E. F., Goddard, T. D., Huang, C. C., Couch, G. S., Greenblatt, D. M., Meng, E. C., & Ferrin, T. E. (2004). UCSF Chimera - A visualization system for exploratory research and analysis. Journal of Computational Chemistry, 25(13), 1605–1612. https://doi.org/10.1002/jcc.20084

Protzer, U., Seyfried, S., Quasdorff, M., Sass, G., Svorcova, M., Webb, D., Bohne, F., Hösel, M., Schirmacher, P., & Tiegs, G. (2007). Antiviral Activity and Hepatoprotection by Heme Oxygenase-1 in Hepatitis B Virus Infection. Gastroenterology, 133(4), 1156–1165. https://doi.org/10.1053/j.gastro.2007.07.021

Ratajczak, M. Z., & Kucia, M. (2020). SARS-CoV-2 infection and overactivation of Nlrp3 inflammasome as a trigger of cytokine “storm” and risk factor for damage of hematopoietic stem cells. Leukemia, 34(7), 1726–1729. https://doi.org/10.1038/s41375-020-0887-9

Rossi, M., Delbauve, S., Wespes, E., Roumeguère, T., Leo, O., Flamand, V., Le Moine, A., & Hougardy, J. M. (2018). Dual effect of hemin on renal ischemia-reperfusion injury. Biochemical and Biophysical Research Communications, 503(4), 2820–2825. https://doi.org/10.1016/j.bbrc.2018.08.046

Rossi, M., Piagnerelli, M., Van Meerhaeghe, A., & Zouaoui Boudjeltia, K. (2020). Heme oxygenase-1 (HO-1) cytoprotective pathway: A potential treatment strategy against coronavirus disease 2019 (COVID-19)-induced cytokine storm syndrome. Medical Hypotheses, 144, 110242. https://doi.org/10.1016/j.mehy.2020.110242

San Juan, I., Bruzzone, C., Bizkarguenaga, M., Bernardo-Seisdedos, G., Laín, A., Gil-Redondo, R., Diercks, T., Gil-Martínez, J., Urquiza, P., Arana, E., Seco, M., García de Vicuña, A., Embade, N., Mato, J. M., & Millet, O. (2020). Abnormal concentration of porphyrins in serum from COVID-19 patients. In British Journal of Haematology (Vol. 190, Issue 5, pp. e265–e267). Blackwell Publishing Ltd. https://doi.org/10.1111/bjh.17060

Singh, D., Wasan, H., & Reeta, K. H. (2020). Heme oxygenase-1 modulation: A potential therapeutic target for COVID-19 and associated complications. Free Radical Biology and Medicine, 161, 263–271. https://doi.org/10.1016/j.freeradbiomed.2020.10.016

Siu, K. L., Yuen, K. S., Castano-Rodriguez, C., Ye, Z. W., Yeung, M. L., Fung, S. Y., Yuan, S., Chan, C. P., Yuen, K. Y., Enjuanes, L., & Jin, D. Y. (2019). Severe acute respiratory syndrome Coronavirus ORF3a protein activates the NLRP3 inflammasome by promoting TRAF3-dependent ubiquitination of ASC. FASEB Journal, 33(8), 8865–8877. https://doi.org/10.1096/fj.201802418R

Su, W. L., Lin, C. P., Hang, H. C., Wu, P. S., Cheng, C. F., & Chao, Y. C. (2021). Desaturation and heme elevation during COVID-19 infection: A potential prognostic factor of heme oxygenase-1. Journal of Microbiology, Immunology and Infection, 54(1), 113–116. https://doi.org/10.1016/j.jmii.2020.10.001

Swanson, K. V., Deng, M., & Ting, J. P. Y. (2019). The NLRP3 inflammasome: molecular activation and regulation to therapeutics. In Nature Reviews Immunology (Vol. 19, Issue 8, pp. 477–489). Nature Publishing Group. https://doi.org/10.1038/s41577-019-0165-0

Takaki, S., Takeyama, N., Kajita, Y., Yabuki, T., Noguchi, H., Miki, Y., Inoue, Y., Nakagawa, T., & Noguchi, H. (2010). Beneficial effects of the heme oxygenase-1/carbon monoxide system in patients with severe sepsis/septic shock. Intensive Care Medicine, 36(1), 42–48. https://doi.org/10.1007/s00134-009-1575-4

Tan, M. E., Li, J., Xu, H. E., Melcher, K., & Yong, E. L. (2015). Androgen receptor: Structure, role in prostate cancer and drug discovery. In Acta Pharmacologica Sinica (Vol. 36, Issue 1, pp. 3–23). Nature Publishing Group. https://doi.org/10.1038/aps.2014.18

Tay, M. Z., Poh, C. M., Rénia, L., MacAry, P. A., & Ng, L. F. P. (2020). The trinity of COVID-19: immunity, inflammation and intervention. In Nature Reviews Immunology (Vol. 20, Issue 6, pp. 363–374). Nature Research. https://doi.org/10.1038/s41577-020-0311-8

Trott, O., & Olson, A. J. (2009). AutoDock Vina: Improving the speed and accuracy of docking with a new scoring function, efficient optimization, and multithreading. Journal of Computational Chemistry, 31(2), NA–NA. https://doi.org/10.1002/jcc.21334

Tseng, C. K., Lin, C. K., Wu, Y. H., Chen, Y. H., Chen, W. C., Young, K. C., & Lee, J. C. (2016). Human heme oxygenase 1 is a potential host cell factor against dengue virus replication. Scientific Reports, 6(1), 1–16. https://doi.org/10.1038/srep32176

van den Berg, D. F., & te Velde, A. A. (2020). Severe COVID-19: NLRP3 Inflammasome Dysregulated. In Frontiers in Immunology (Vol. 11, p. 1580). Frontiers Media S.A. https://doi.org/10.3389/fimmu.2020.01580

Vitali, S. H., Fernandez-Gonzalez, A., Nadkarni, J., Kwong, A., Rose, C., Alex Mitsialis, S., & Kourembanas, S. (2020). Heme oxygenase-1 dampens the macrophage sterile inflammasome response and regulates its components in the hypoxic lung. American Journal of Physiology - Lung Cellular and Molecular Physiology, 318(1), L125–L134. https://doi.org/10.1152/ajplung.00074.2019

Wagener, F. A. D. T. G., Pickkers, P., Peterson, S. J., Immenschuh, S., & Abraham, N. G. (2020). Targeting the heme-heme oxygenase system to prevent severe complications following covid-19 infections. Antioxidants, 9(6), 1–11. https://doi.org/10.3390/antiox9060540

Wang, C., Zhang, Y., Han, L. L., Guo, L., Zhong, H., & Wang, J. (2017). Hemin ameliorates influenza pneumonia by attenuating lung injury and regulating the immune response. International Journal of Antimicrobial Agents, 49(1), 45–52. https://doi.org/10.1016/j.ijantimicag.2016.09.030

Wen, T., Wu, Z. M., Liu, Y., Tan, Y. F., Ren, F., & Wu, H. (2007). Upregulation of heme oxygenase-1 with hemin prevents d-galactosamine and lipopolysaccharide-induced acute hepatic injury in rats. Toxicology, 237(1-3), 184–193. https://doi.org/10.1016/j.tox.2007.05.014

Wu, B., Wu, Y., & Tang, W. (2019). Heme Catabolic Pathway in Inflammation and Immune Disorders. Frontiers in Pharmacology, 10, 825. https://doi.org/10.3389/fphar.2019.00825

Wu, C., Lu, W., Zhang, Y., Zhang, G., Shi, X., Hisada, Y., Grover, S. P., Zhang, X., Li, L., Xiang, B., Shi, J., Li, X. A., Daugherty, A., Smyth, S. S., Kirchhofer, D., Shiroishi, T., Shao, F., Mackman, N., Wei, Y., & Li, Z. (2019). Inflammasome Activation Triggers Blood Clotting and Host Death through Pyroptosis. Immunity, 50(6), 1401–1411.e4. https://doi.org/10.1016/j.immuni.2019.04.003

Xue, H., Guo, H., Li, Y. C., & Hao, Z. M. (2007). Heme oxygenase-1 induction by hemin protects liver cells from ischemia/reperfusion injury in cirrhotic rats. World Journal of Gastroenterology, 13(40), 5384–5390. https://doi.org/10.3748/wjg.v13.i40.5384

Yağci, S., Serin, E., Acicbe, Ö., Zeren, M. İ., & Odabaşı, M. S. (2021). The relationship between serum erythropoietin, hepcidin, and haptoglobin levels with disease severity and other biochemical values in patients with COVID-19. International Journal of Laboratory Hematology, 00, 1–10. https://doi.org/10.1111/ijlh.13479

Yamauchi, T., Lin, Y., Sharp, F. R., & Noble-Haeusslein, L. J. (2004). Hemin induces heme oxygenase-1 in spinal cord vasculature and attenuates barrier disruption and neutrophil infiltration in the injured murine spinal cord. Journal of Neurotrauma, 21(8), 1017–1030. https://doi.org/10.1089/0897715041651042

Yoshihara, E., Masaki, S., Matsuo, Y., Chen, Z., Tian, H., & Yodoi, J. (2013). Thioredoxin/Txnip: Redoxisome, as a redox switch for the pathogenesis of diseases. In Frontiers in Immunology (Vol. 4, Issue DEC, p. 514). Frontiers Media S.A. https://doi.org/10.3389/fimmu.2013.00514

Yu, X., Han, W., Wang, C., Sui, D., Bian, J., Bo, L., & Deng, X. (2018). Upregulation of heme oxygenase-1 by hemin alleviates sepsis-induced muscle wasting in mice. Oxidative Medicine and Cellular Longevity, 2018. https://doi.org/10.1155/2018/8927104

Yuan, Y., Pei, J., & Lai, L. (2013). Binding Site Detection and Druggability Prediction of Protein Targets for Structure-Based Drug Design. Current Pharmaceutical Design, 19(12), 2326–2333. https://doi.org/10.2174/1381612811319120019

Zhu, Z., Wilson, A. T., Mathahs, M., Wen, F., Brown, K. E., Luxon, B. A., & Schmidt, W. N. (2008). Heme oxygenase-1 suppresses hepatitis C virus replication and increases resistance of hepatocytes to oxidant injury. Hepatology, 48(5), 1430–1439. https://doi.org/10.1002/hep.22491

