## Supplemental Raw Data for "Hemin shows antiviral activity *in vitro*, possibly through suppression of viral entry mediators"

| 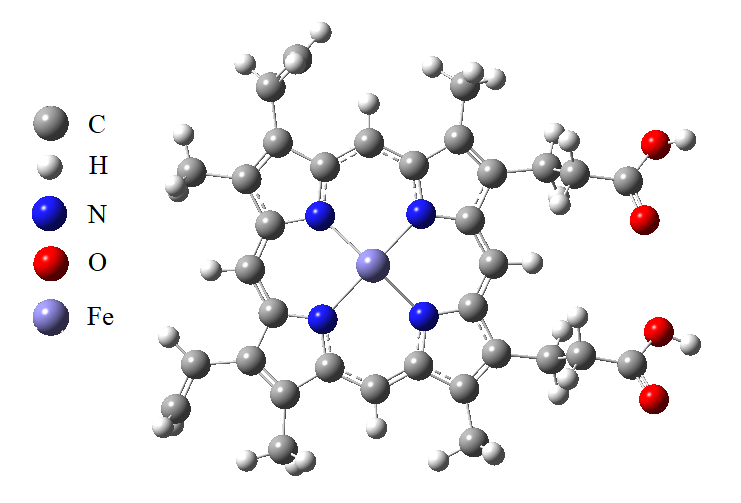  A | 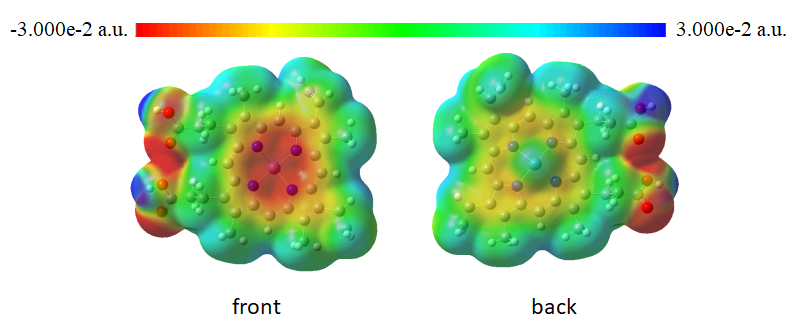  B |
| --- | --- |
| **Figure S1 –** A) Minimized structure and B) electrostatic potential (ESP) map of Hemin. Calculated electronic energy (E)= -3098.198461 Hartree (-84306.267942 eV) and dipole moment (μ)=3.235 Debye. | |

| 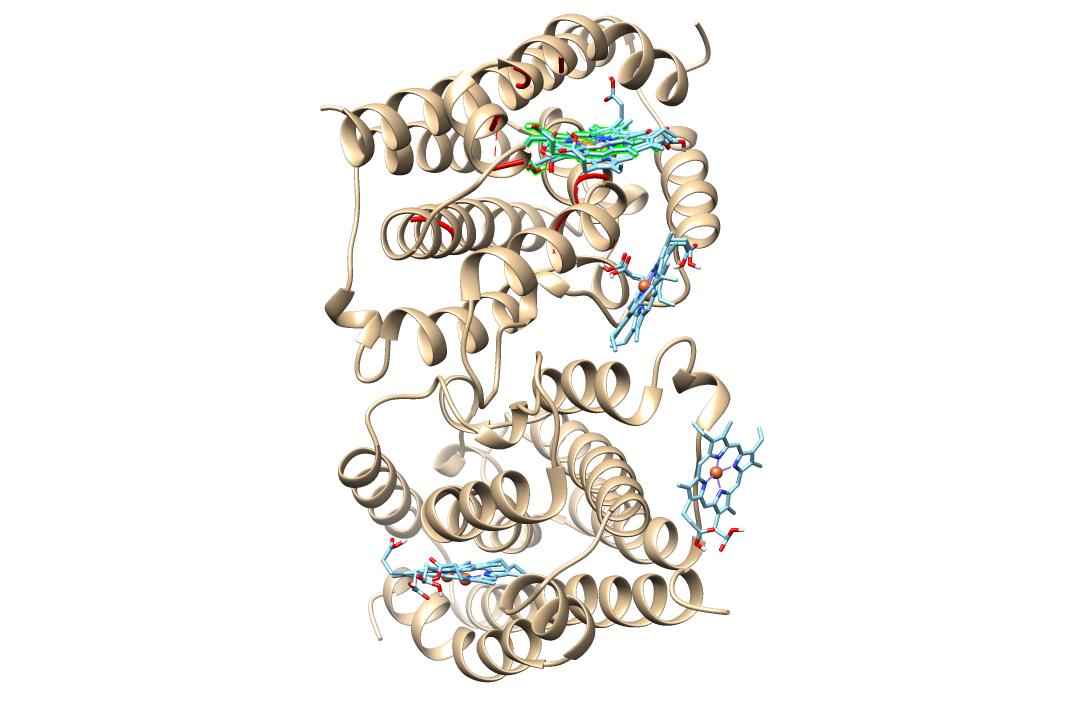  A | 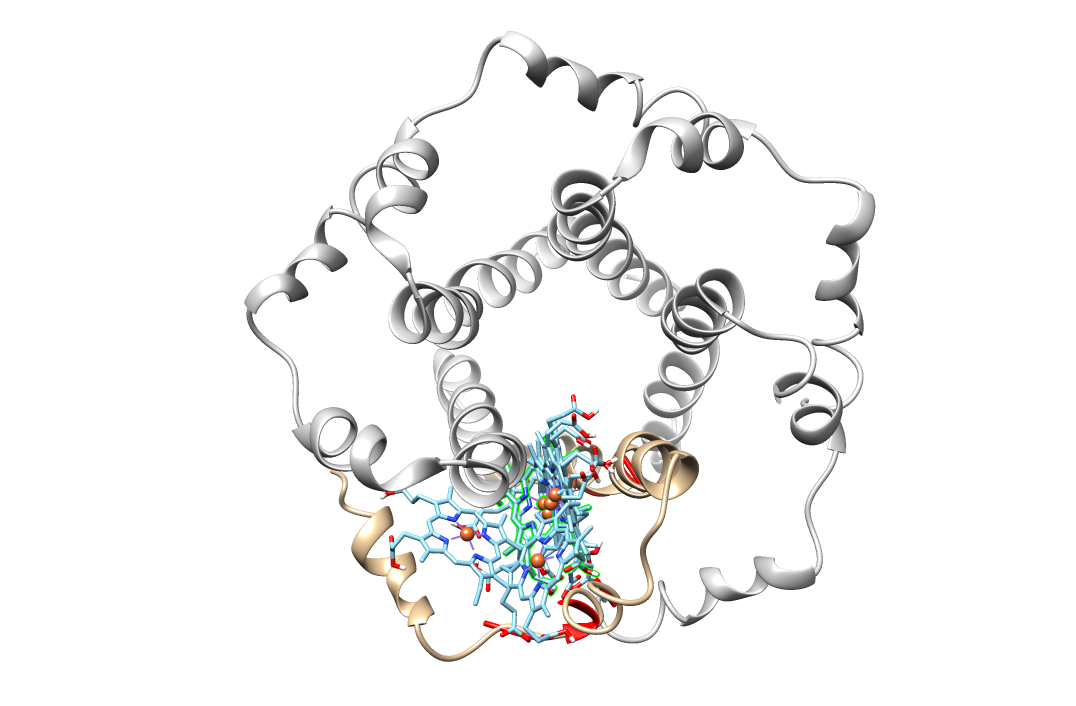  B |
| --- | --- |
| mode \| affinity \| dist from best mode  \| (kcal/mol) \| rmsd l.b.\| rmsd u.b.  -----+------------+----------+----------  1 -9.2 0.000 0.000  2 -9.0 4.168 9.674  3 -8.1 43.044 44.526  4 -7.8 13.858 17.372  5 -7.8 2.102 2.974  6 -7.7 4.100 9.493  7 -7.4 33.540 36.084  8 -7.3 13.788 16.790  9 -7.3 42.714 44.479 | mode \| affinity \| dist from best mode  \| (kcal/mol) \| rmsd l.b.\| rmsd u.b.  -----+------------+----------+----------  1 -6.0 0.000 0.000  2 -5.9 2.602 6.063  3 -5.9 2.361 3.853  4 -5.8 5.538 9.005  5 -5.7 5.829 9.463  6 -5.7 2.789 6.644  7 -5.6 9.588 12.337  8 -5.6 5.529 9.711  9 -5.5 11.740 15.584 |
| **Figure S2 –** A) Hemin-6EHA B) Hemin-5X29 complexes with 9 highest affinity energy conformations, The red colored amino acids indicate the highest energy druggabale cavity (druggability score, 6EHA= 1878; 5X29= 4232) and the highest energy conformation is green colored | |

| 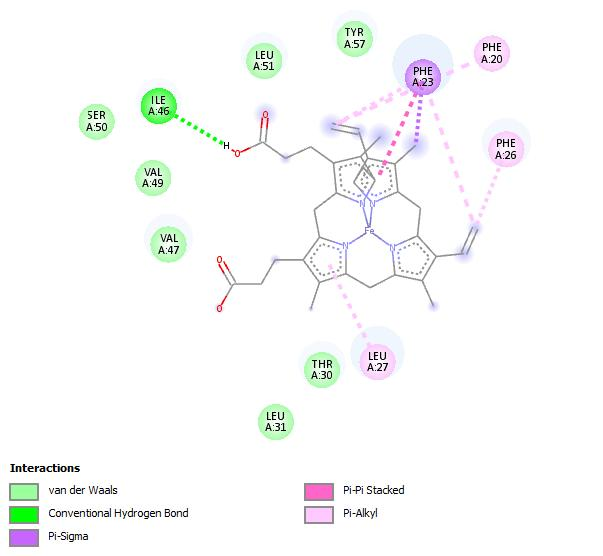  mode 1 | 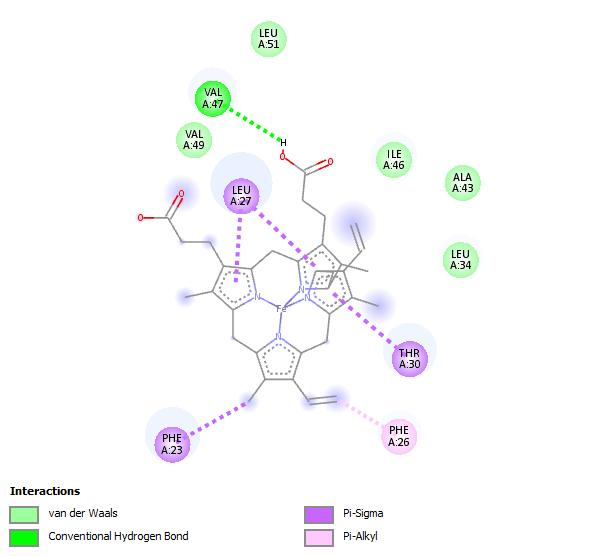  mode 2 |
| --- | --- |
| 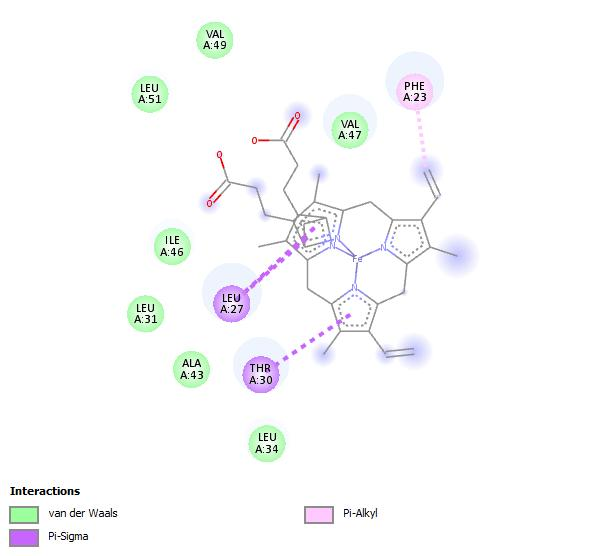  mode 3 | 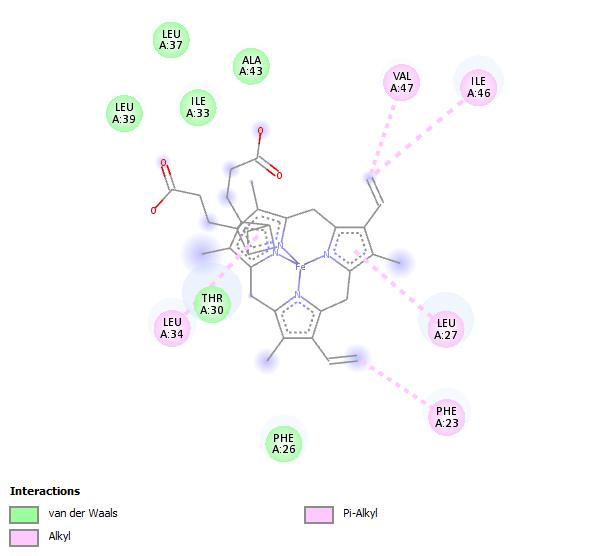  mode 4 |
| 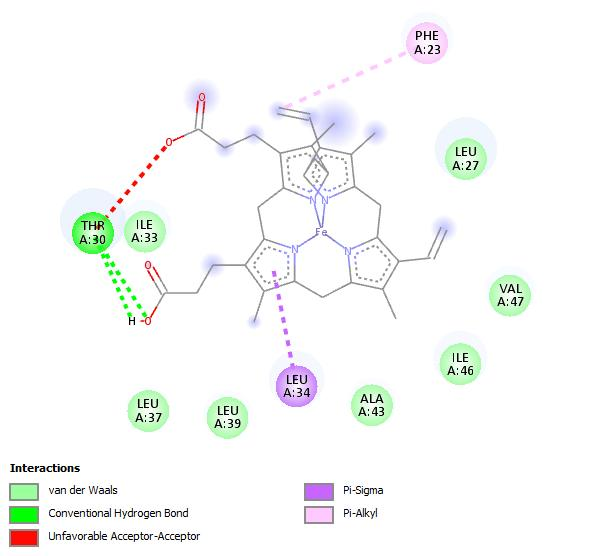  mode 5 | 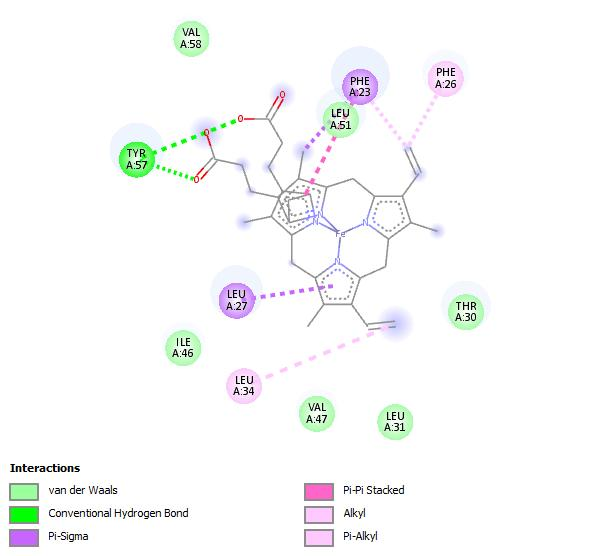  mode 6 |
| 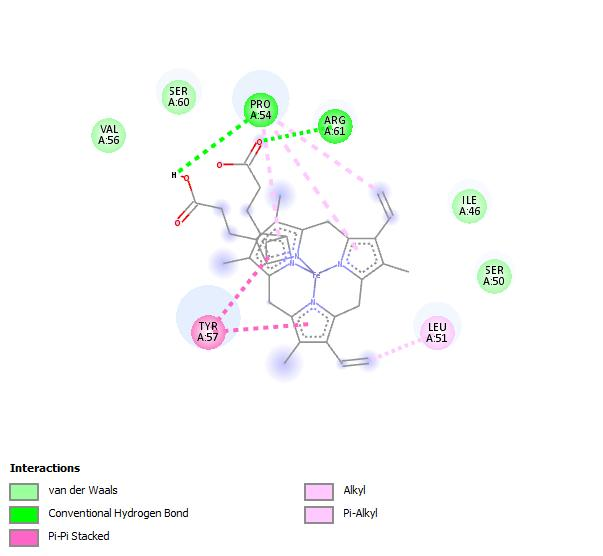  mode 7 | 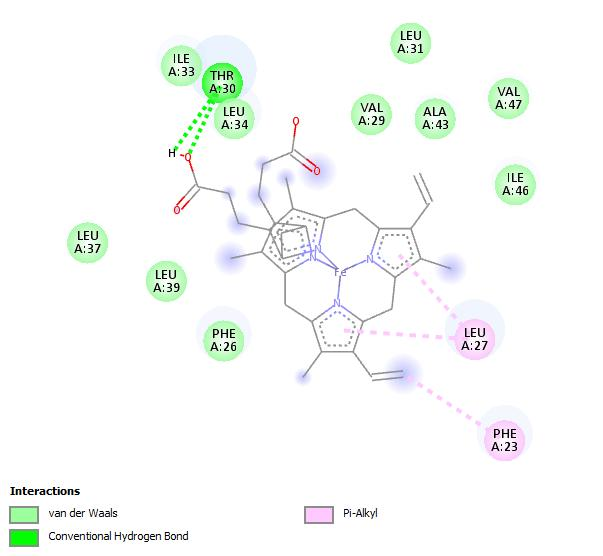  mode 8 |
| 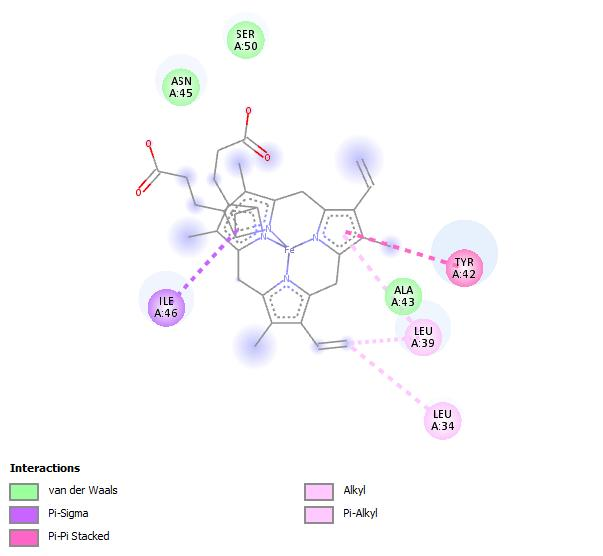  mode 9 | 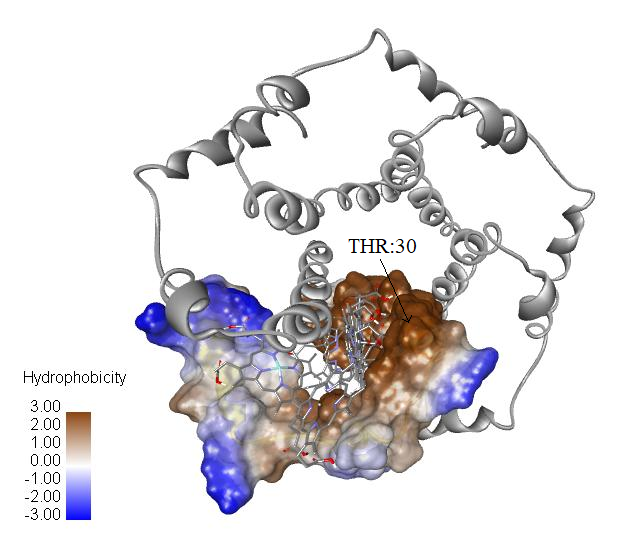  A |
| **Figure S3 –** 2D interaction maps of hemin-5X29 complex for all conformations and A) hydrophobic property | |

| 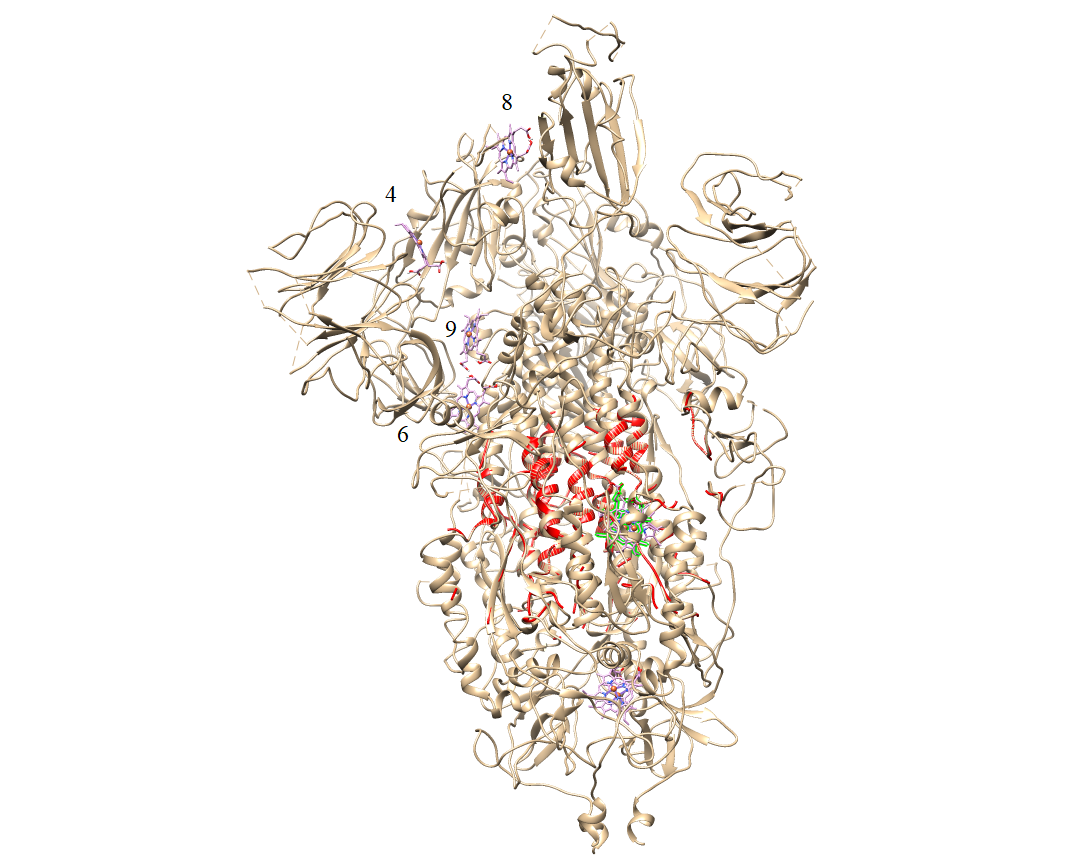  6VSB  a | 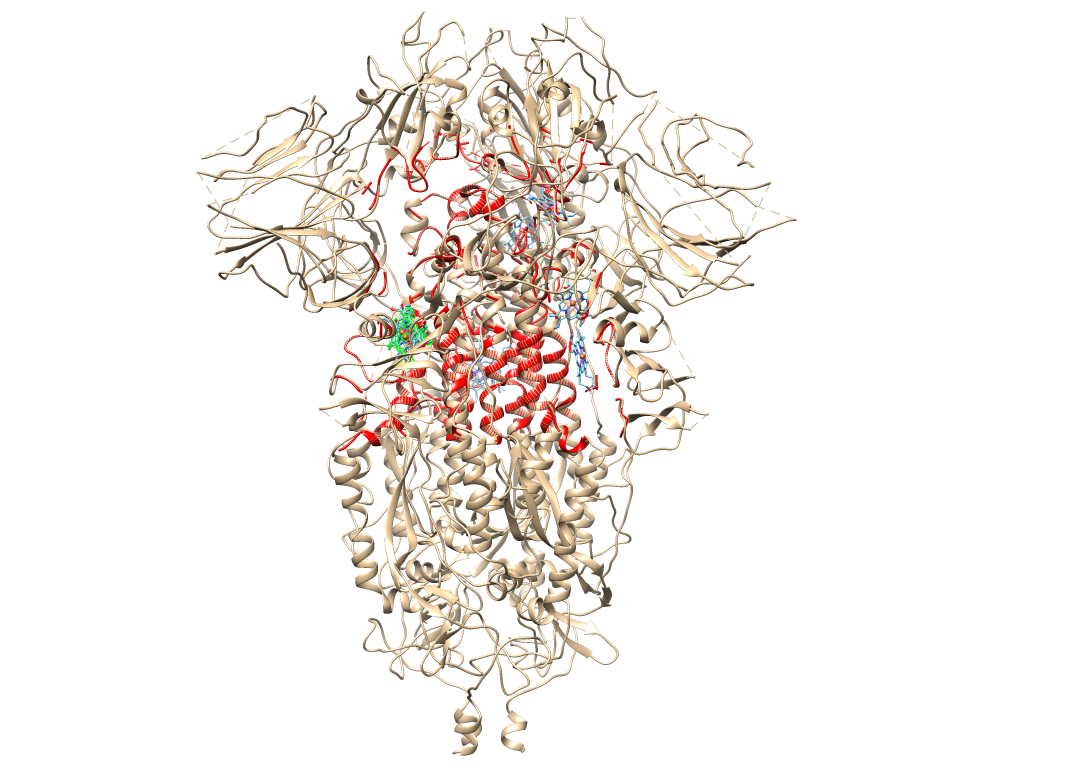  6VXX  b |
| --- | --- |
| mode \| affinity \| dist from best mode  \| (kcal/mol) \| rmsd l.b.\| rmsd u.b.  -----+------------+----------+----------  1 -8.7 0.000 0.000  2 -8.6 31.119 34.523  3 -8.2 2.110 4.834  4 -8.0 66.635 71.861  5 -7.9 30.933 35.296  6 -7.9 37.125 41.480  7 -7.9 2.406 4.056  8 -7.6 75.207 80.578  9 -7.5 46.627 51.814 | mode \| affinity \| dist from best mode  \| (kcal/mol) \| rmsd l.b.\| rmsd u.b.  -----+------------+----------+----------  1 -8.8 0.000 0.000  2 -8.3 1.621 2.092  3 -8.0 1.041 5.826  4 -7.8 32.173 36.452  5 -7.7 35.241 37.634  6 -7.5 34.289 36.703  7 -7.5 38.169 40.196  8 -7.3 33.434 35.751  9 -7.2 39.622 43.658 |
| 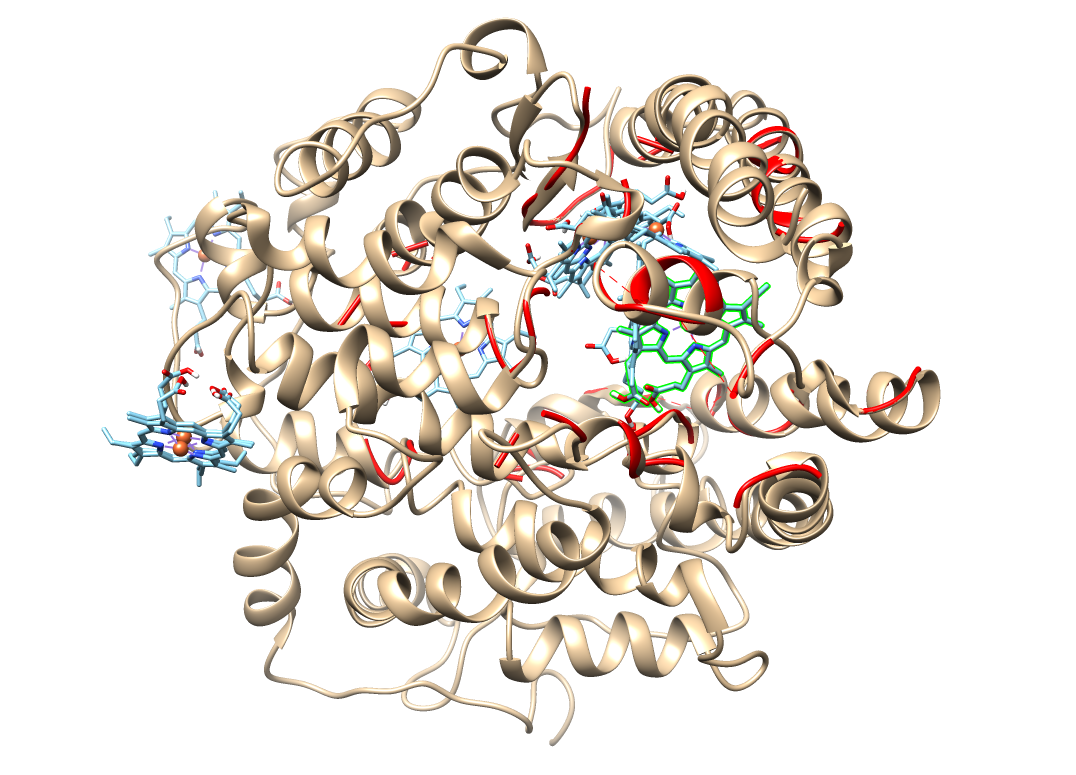  1R42  c | 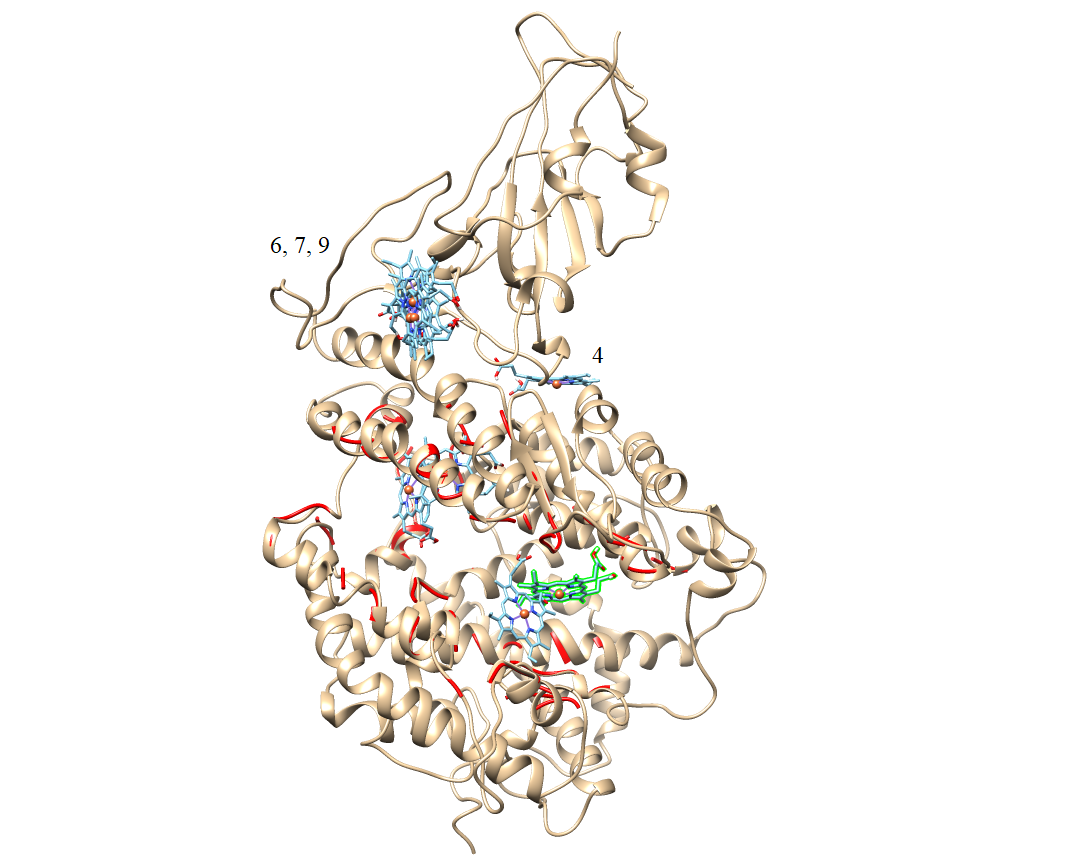  6M0J  d |
| mode \| affinity \| dist from best mode  \| (kcal/mol) \| rmsd l.b.\| rmsd u.b.  -----+------------+----------+----------  1 -7.9 0.000 0.000  2 -7.8 4.893 6.412  3 -7.6 13.971 16.808  4 -7.5 36.534 41.050  5 -7.4 9.109 10.573  6 -7.4 22.223 25.562  7 -7.2 15.048 15.842  8 -7.1 36.307 41.202  9 -7.1 46.510 49.663 | mode \| affinity \| dist from best mode  \| (kcal/mol) \| rmsd l.b.\| rmsd u.b.  -----+------------+----------+----------  1 -8.0 0.000 0.000  2 -7.6 20.158 23.986  3 -7.5 6.636 9.447  4 -7.4 27.406 29.200  5 -7.4 17.718 21.590  6 -7.4 39.909 43.885  7 -7.3 39.632 44.009  8 -7.2 42.602 46.165  9 -7.1 40.900 45.284 |
| **Figure S4A –**The effect of the nine highest affinity hemin conformations on different regions of 6VSB, 6VX, 1R42 and 6M0J proteins (supplementary, DockingRawData.zip), (druggability score, 6VSB= 10509; 6VXX= 16603; 1R42= 6822 and 6M0J= 7414). Gold color: protein, red: druggable active area of protein with the highest score, and green: conformation with the highest affinity.  According to UniProtKB database (https://www.uniprot.org/uniprot/P0DTC2), 6VYB (open form) and 6VXX (close form) spike proteins have two binding areas. These are amino acids between 319-541 (Receptor Binding Domain, RBD) and 437-508 (receptor-binding motif). Besides, 30-41, 82-84 and 353-357 amino acids are active in binding of ACE2 protein (1R42) to spike proteins (https://www.uniprot.org/uniprot/Q9BYF1). When the effect of hemin against these proteins and Spike-ACE2 bounded protein complex is examined (Figure S4A), it can be said that it has good activity (from -8.0 to -7.1 kcal/mol). | |

| 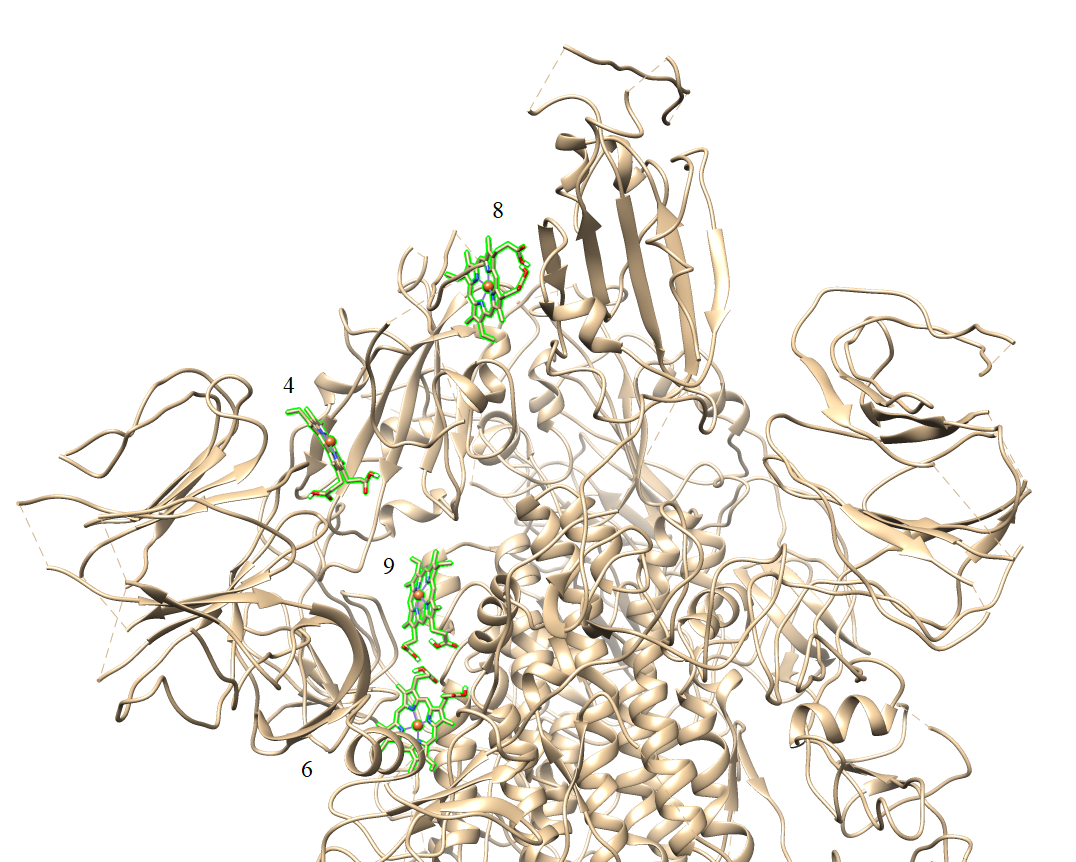  6VSB with 4, 6, 8 and 9 conformations of Hemin  e |  |
| --- | --- |
| 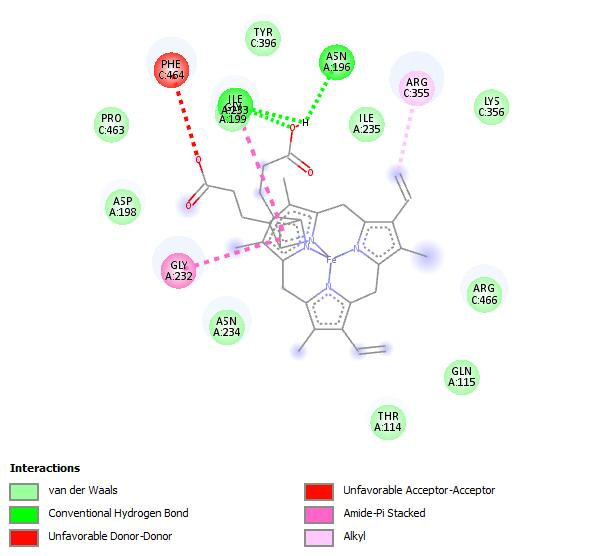  6VSB - 4^th^ conformation  f | 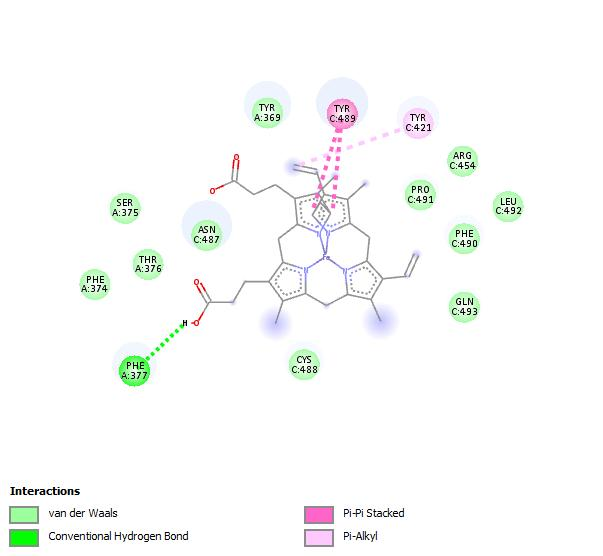  6VSB - 8^th^ conformation  g |
| 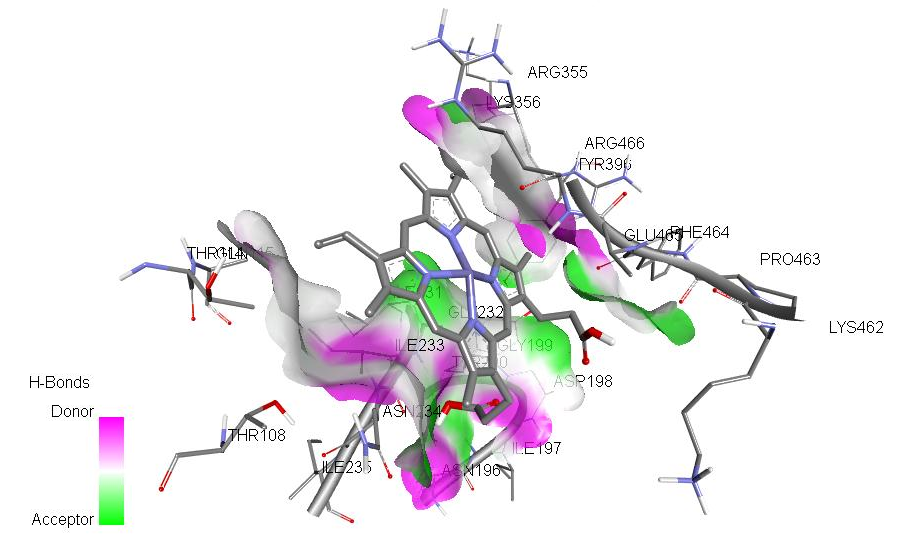  6VSB - 4^th^ conformation, 3D H-bond pose  h | 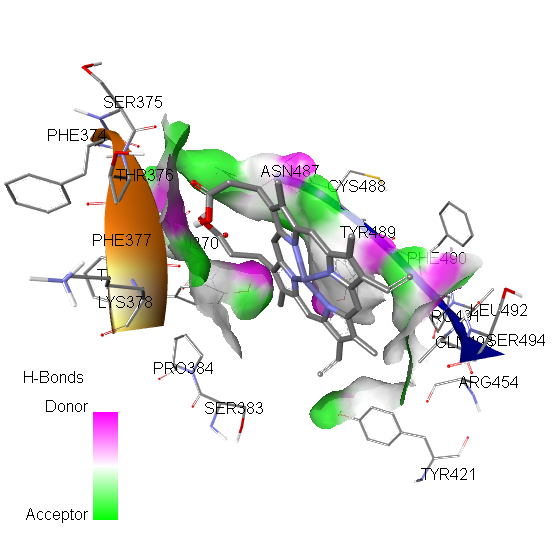  6VSB - 8^th^ conformation, 3D H-bond pose  i |
| 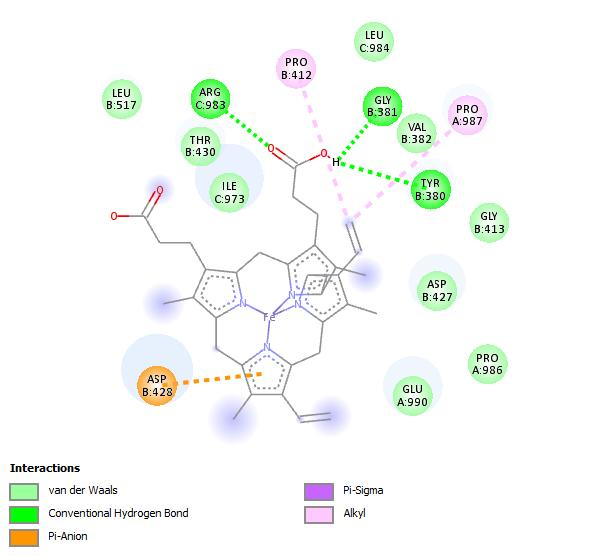  6VXX - 7^th^ conformation  j | 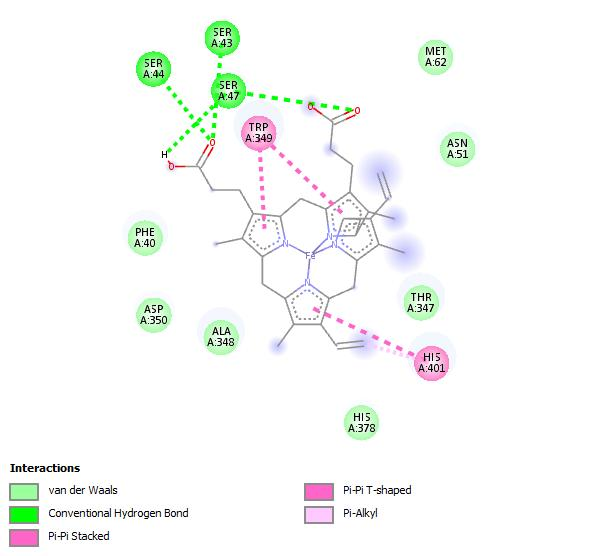  1R42 - 3^rt^ conformation  k |
| 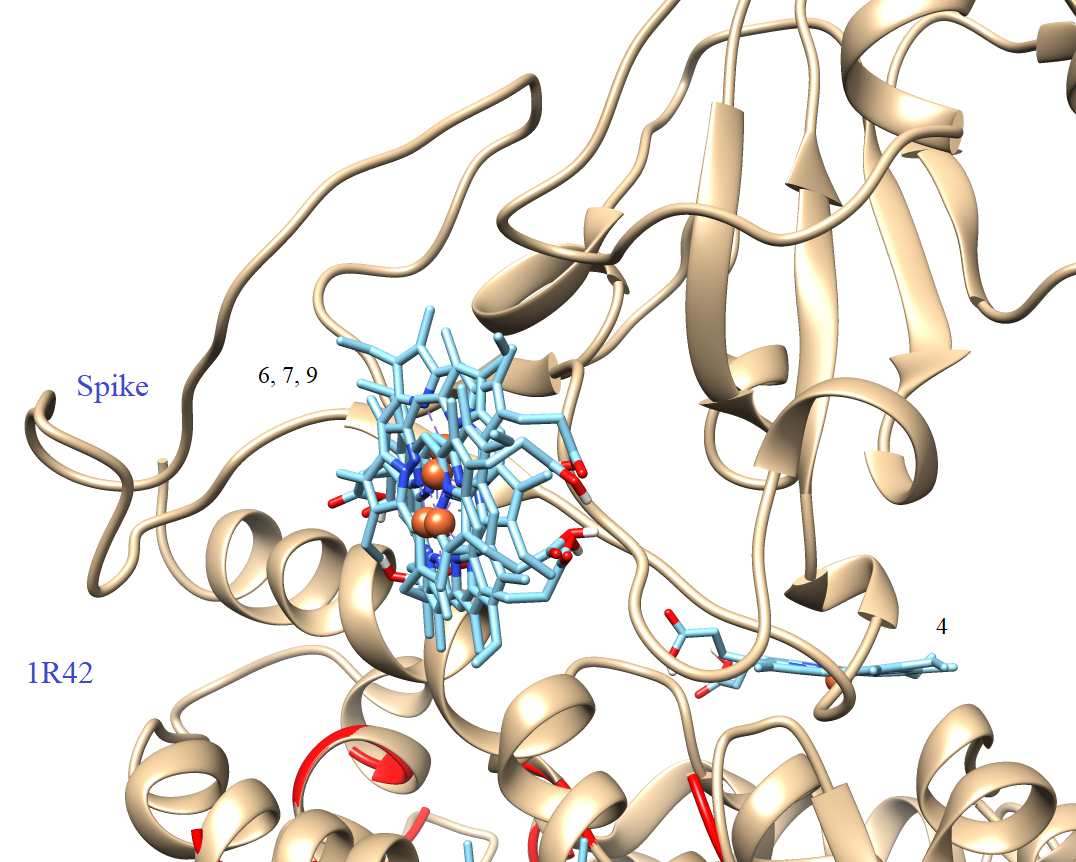  6M0J (Spike-ACE2 bounded) with 4, 6, 7, and 9 conformations of Hemin  l |  |
| 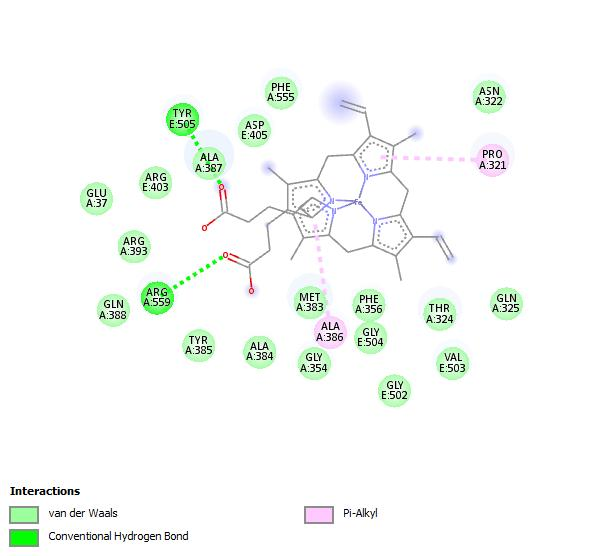  6M0J - 4^th^ conformation  m | 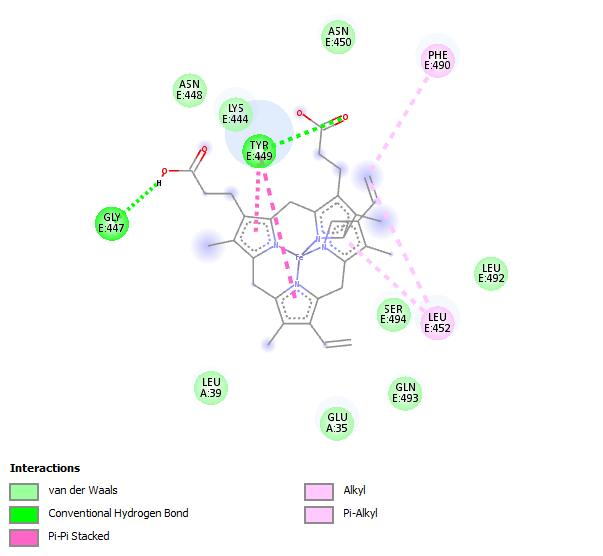  6M0J - 6^th^ conformation  n |
| 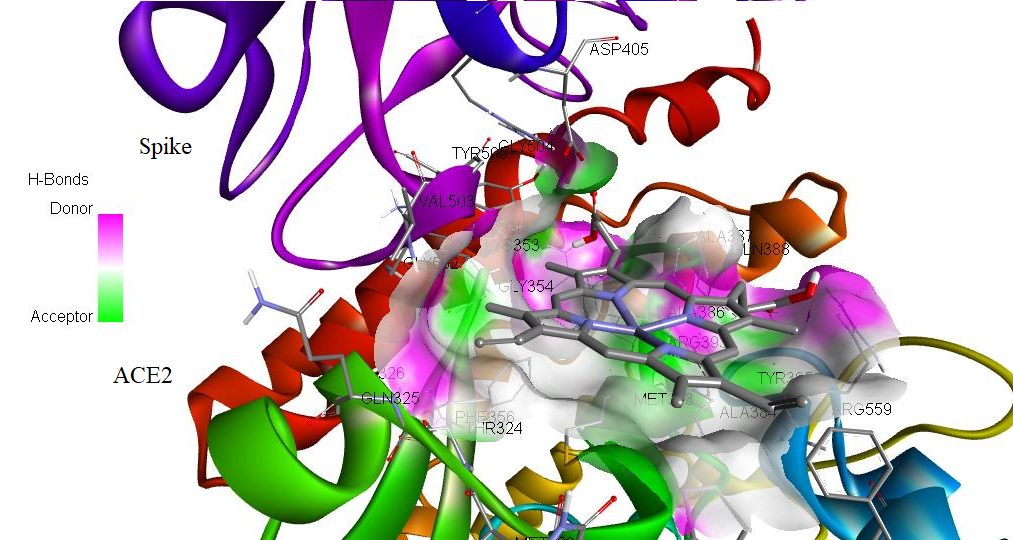  6M0J - 4^th^ conformation, 3D H-bond pose  o | 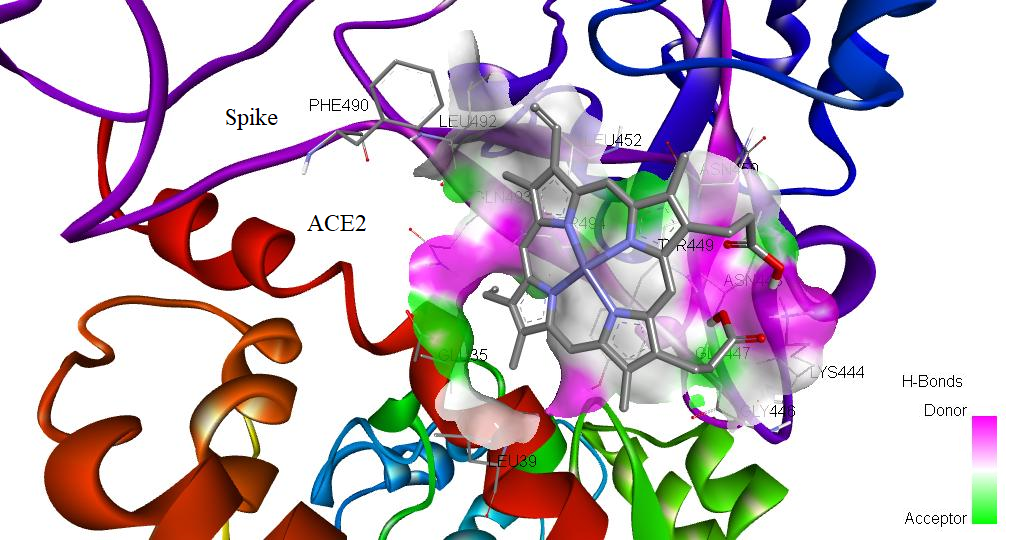  6M0J - 6^th^ conformation, 3D H-bond pose  p |
| 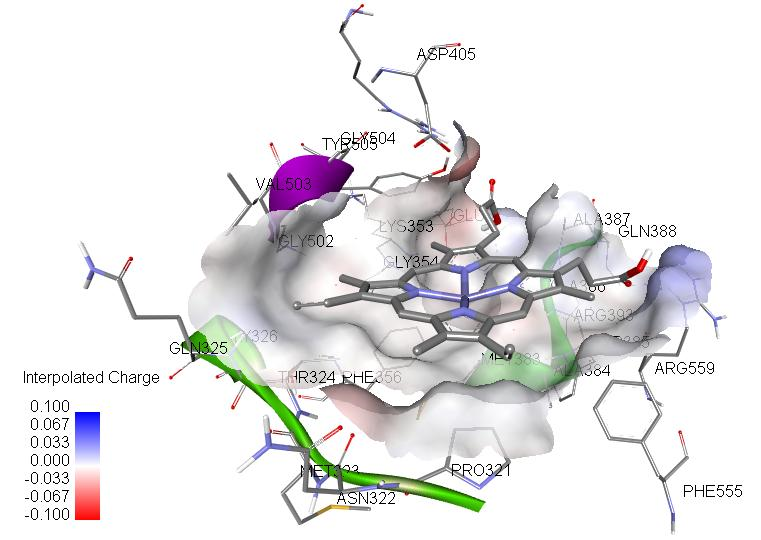  6M0J - 4^th^ conformation, 3D interpolated charge pose  q | 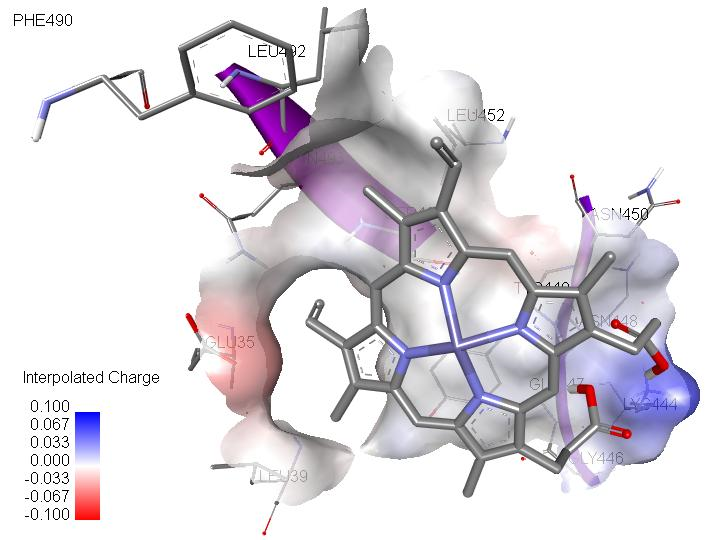  6M0J - 4^th^ conformation, 3D interpolated charge pose  r |
| 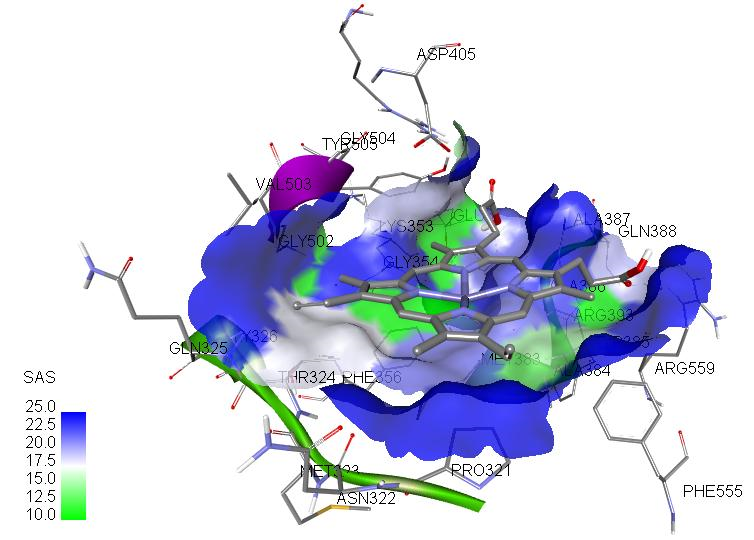  6M0J - 4^th^ conformation, 3D interpolated charge pose  s | 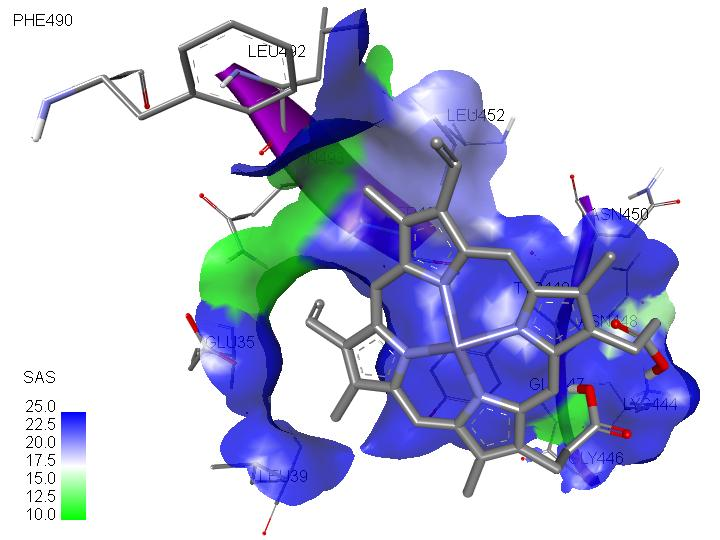  6M0J - 4^th^ conformation, 3D interpolated charge pose  t |
| 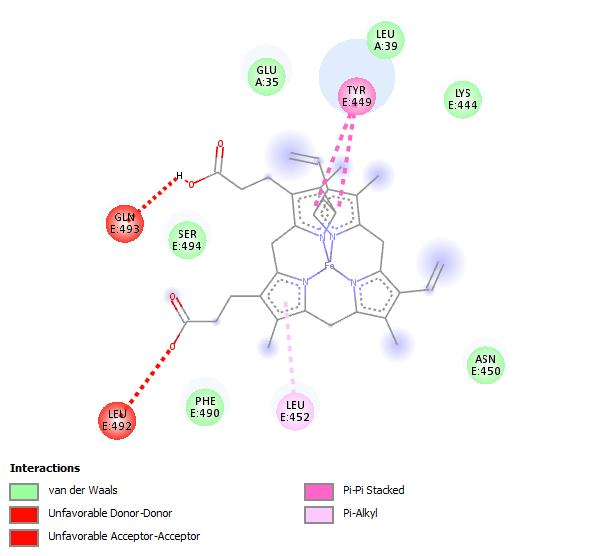  6M0J - 7^th^ conformation  u | 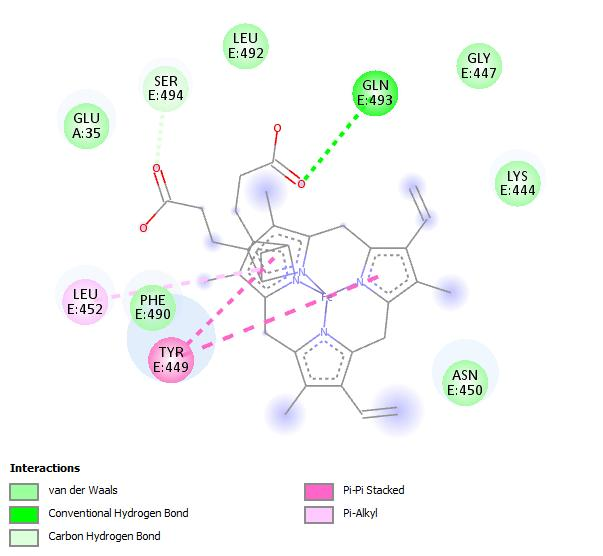  6M0J - 9^th^ conformation  v |
| 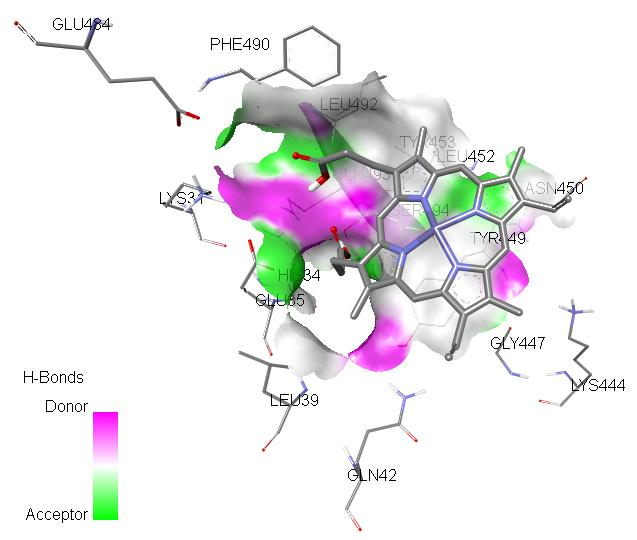  6M0J - 7^th^ conformation, 3D H-bond pose  w | 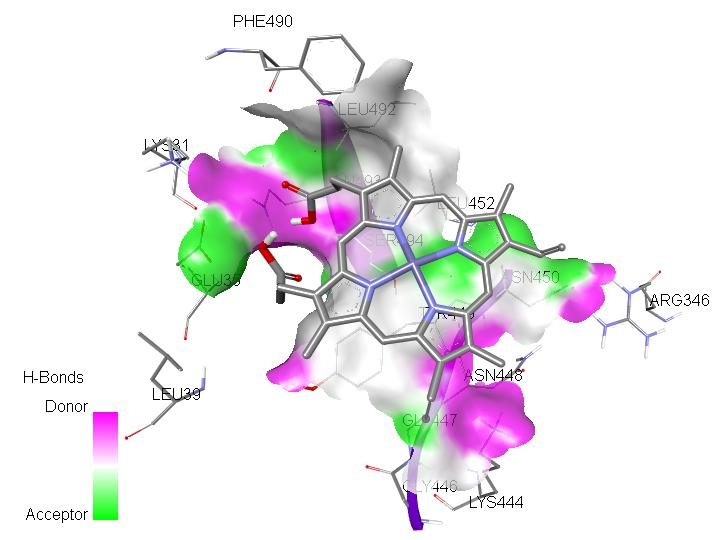  6M0J - 9^th^ conformation, 3D H-bond pose  x |
| **Figure S4B –** 3D and 2D representations of the effect of important Hemin conformations on the 6VSB, 6VX, 1R42 and 6M0J protein regions  **S4Be-k:** Two-dimensional interaction maps show amino acids that these two conformations can affect at the binding site of spike proteins: 6VSB; (4th) ARG:355, LYS:356, TYR:396, PRO:463, PHE:464, ARG:466; (8th) PHE:374, TYR:369, SER:375, THR:376, PHE:377, TYR:421, ARG:454, ASN:487, CYS:488, TYR:489, PHE:490, PRO:491 , LEU:492,GLN493 (RBD region amino acids).  **S4Bm-x:** 1R42 (A); GLU:35, LEU:39, GLU:37, GLY:354, PHE:356, and Spike (B); LYS:444, GLY:447, ASN:448, TRY:449, ASN:450, LEU:452, PHE:490, LEU:492, GLN:493, SER:494, GLY:502, VAL:503, GLY504, TRY:505. | |

| 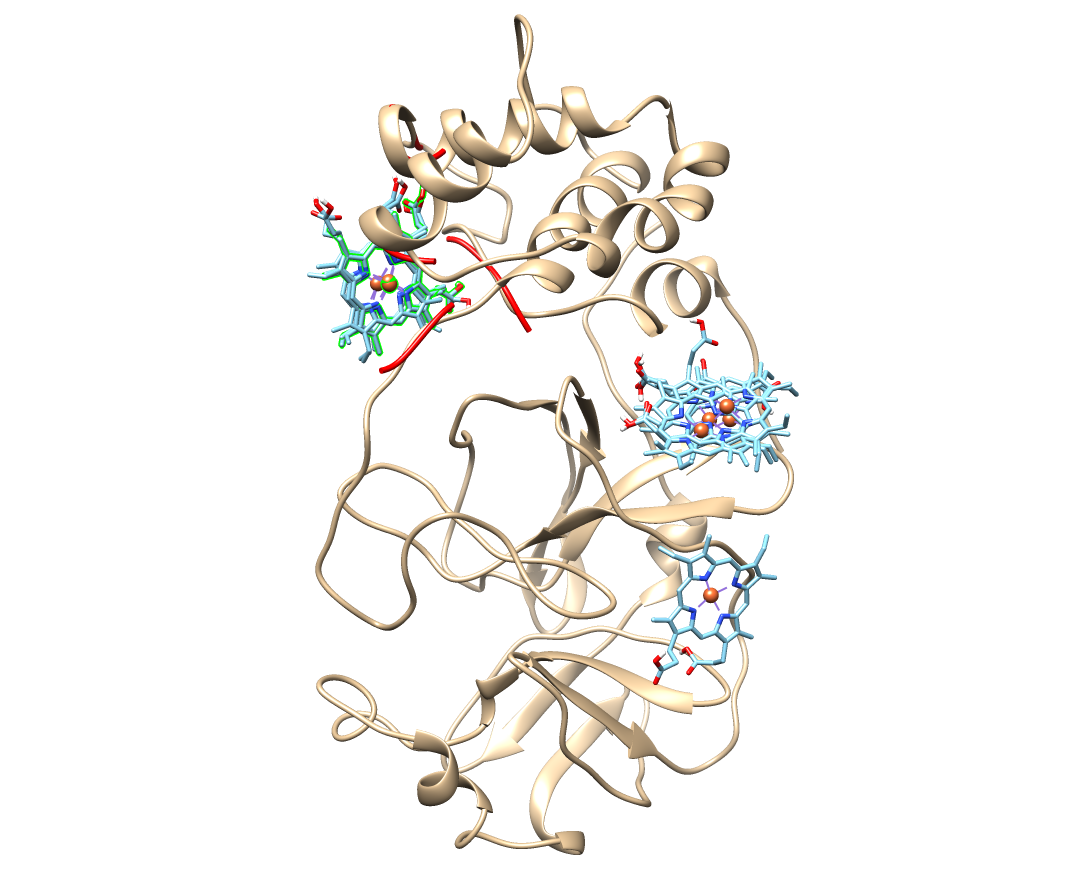  6M03  a | 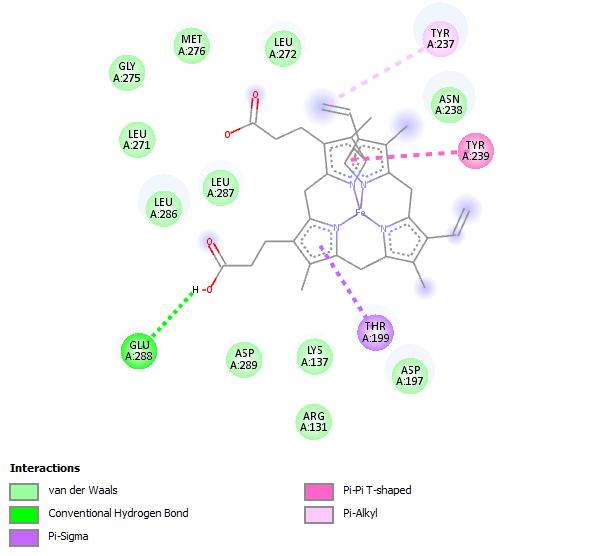  b |
| --- | --- |
| mode \| affinity \| dist from best mode  \| (kcal/mol) \| rmsd l.b.\| rmsd u.b.  -----+------------+----------+----------  1 -8.2 0.000 0.000  2 -8.2 25.261 29.650  3 -8.1 24.220 28.311  4 -8.0 25.172 29.772  5 -7.7 0.739 6.488  6 -7.5 2.462 7.164  7 -7.4 2.192 6.629  8 -7.4 24.945 29.037  9 -7.4 33.729 38.121 |  |
| 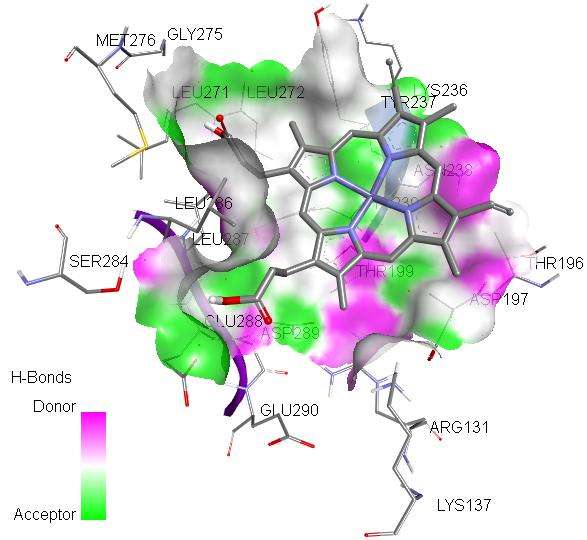  c | 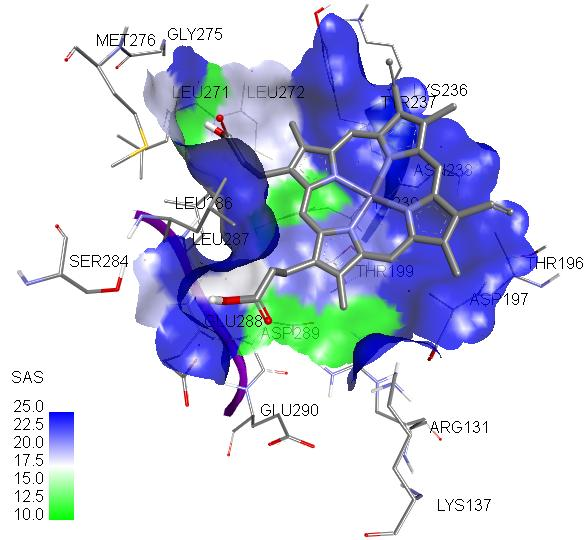  d |
| 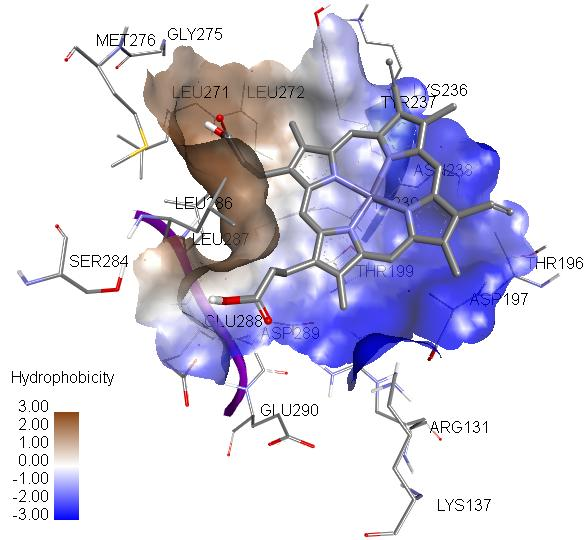  e | 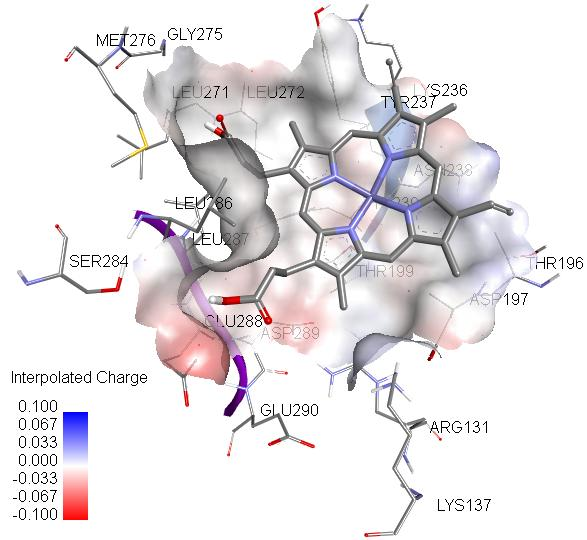  f |
| **Figure S5** – Hemin-6M03 interaction: a) all conformations of hemin (druggability score= 51); interaction of 6M03 with highest energy conformation of hemin: b) two dimentional, c) hydrogen donor-acceptor, d) SAS and e) hydrophobicity and interpolated charge maps | |

| 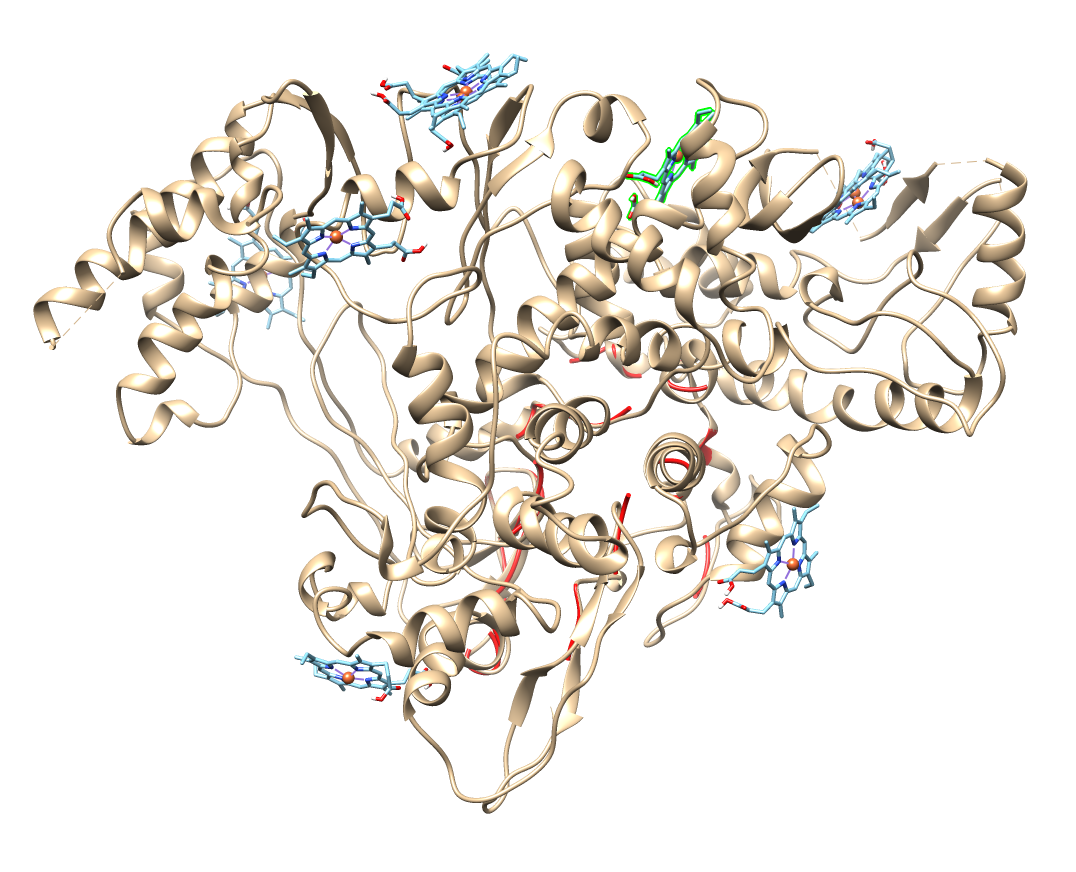  a | 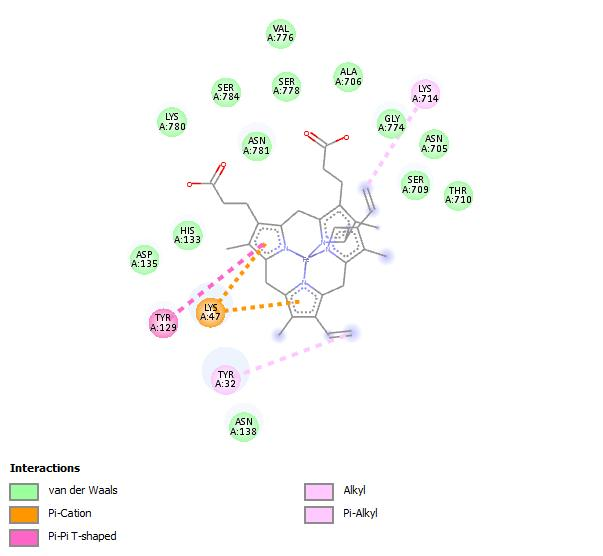  b |
| --- | --- |
| mode \| affinity \| dist from best mode  \| (kcal/mol) \| rmsd l.b.\| rmsd u.b.  -----+------------+----------+----------  1 -7.8 0.000 0.000  2 -7.7 17.403 21.015  3 -7.5 35.947 42.406  4 -7.4 27.223 30.788  5 -7.4 17.317 21.228  6 -7.3 40.089 43.294  7 -7.2 59.003 61.790  8 -7.2 43.171 47.764  9 -7.2 27.221 30.728 |  |
| 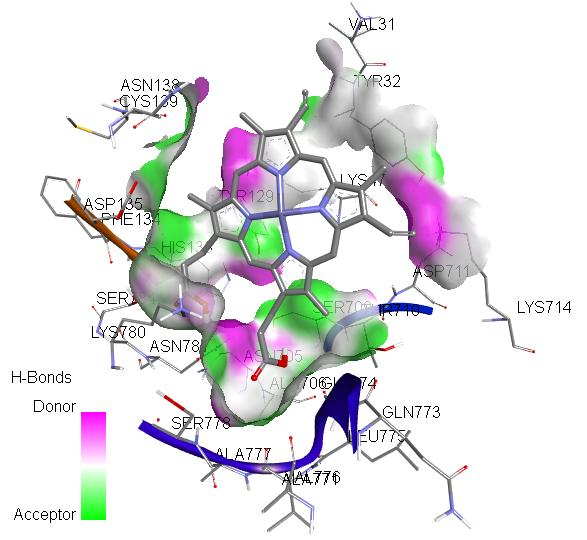  c | 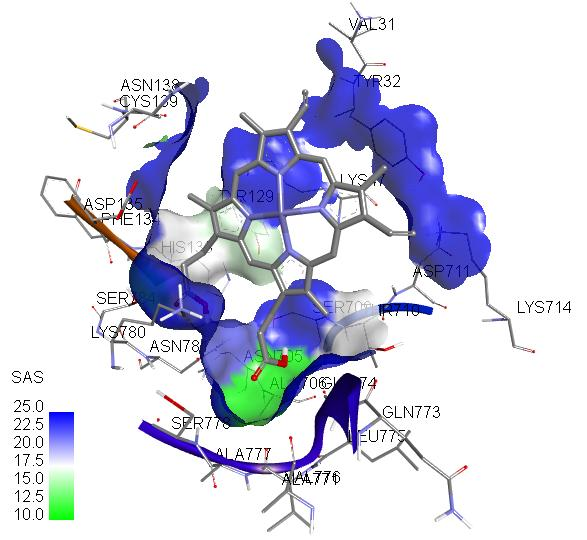  d |
| **Figure S6** - Hemin-6M71 interaction: a) all conformations of hemin (druggability score= 4115); interaction of 6M71 with highest energy conformation of hemin: b) two dimentional, c) hydrogen donor-acceptor and d) SAS maps | |
